# SMC and the bactofilin/PadC scaffold have distinct yet redundant functions in chromosome segregation and organization in *Myxococcus xanthus*

**DOI:** 10.1101/2020.06.17.156356

**Authors:** Deepak Anand, Dominik Schumacher, Lotte Søgaard-Andersen

**Affiliations:** Department of Ecophysiology, Max Planck Institute for Terrestrial Microbiology, Karl-von-Frisch Straße 10, 35043 Marburg, Germany

**Keywords:** Smc, ScpAB, bactofilin, PadC, ParA, ParB

## Abstract

In bacteria, ParAB*S* systems and structural maintenance of chromosome (SMC) condensin-like complexes are important for chromosome segregation and organization. The rod-shaped *Myxococcus xanthus* cells have a unique chromosome arrangement in which a scaffold composed of three bactofilins (BacNOP) and PadC positions the essential ParB·*parS* segregation complexes and the DNA segregation ATPase ParA in the subpolar regions. Here, we identify the Smc and ScpAB subunits of the SMC complex in *M. xanthus* and demonstrate that SMC is conditionally essential with mutants containing *smc* or *scpAB* deletions being temperature sensitive. Lack of SMC caused defects in chromosome segregation and organization. Lack of the BacNOP/PadC scaffold caused chromosome segregation defects but was not essential. Inactivation of SMC was synthetic lethal with lack of the BacNOP/PadC scaffold. Lack of SMC interfered with formation of the BacNOP/PadC scaffold while lack of this scaffold did not interfere with chromosome association by SMC. Altogether, our data support that three systems cooperate to enable chromosome segregation in *M. xanthus*, whereby ParAB*S* constitutes the basic machinery and SMC and the BacNOP/PadC scaffold have distinct yet redundant roles in this process with SMC supporting individualization of daughter chromosomes and BacNOP/PadC making the ParABS system operate more robustly

## Introduction

Chromosome segregation is closely coordinated with cell division to ensure that daughter cells inherit the correct chromosome complement. Bacterial chromosomes are highly compacted and organized to fit into the confines of cells while still allowing DNA replication, repair, recombination and segregation to occur (Badrinarayanan *et al.*, 2015, Dame *et al.*, 2020). This organization includes positioning of individual chromosomal loci to the same subcellular locations in each cell cycle and is established during segregation (Viollier *et al.*, 2004). Because replication and segregation occur in parallel in bacteria, cells face the task of not only faithfully segregating daughter chromosomes to opposite cell halves but simultaneously lay down these chromosomes in a spatially organized manner.

Generally, bacteria contain a single chromosome, replication initiates at a single, well-defined origin of replication (*ori*), proceeds bidirectionally and terminates in the terminus region (*ter*) (Badrinarayanan *et al.*, 2015). The subcellular locations of *ori* and *ter* define two overall chromosome arrangements (Badrinarayanan *et al.*, 2015). In the more frequently described *ori-ter* longitudinal pattern, and as observed for instance in *Caulobacter crescentus* (Viollier *et al.*, 2004), these two regions localize at opposite cell poles, and with the two replichores arranged in close contact in-between and in the same order as the genetic map (Le *et al.*, 2013, Marbouty *et al.*, 2015, Wang *et al.*, 2015, Viollier *et al.*, 2004). During chromosome segregation, one *ori* copy remains in the polar region while the second copy is segregated to the opposite pole. Alternatively, and as described in *Escherichia coli* (Nielsen *et al.*, 2006, Wang *et al.*, 2006, Niki *et al.*, 2000), *ori* and *ter* are localized around midcell and the two replichores are in opposite cell halves. During segregation, the two *ori* copies are segregated symmetrically to opposite cell halves. Two systems have key functions in chromosome organization and segregation in bacteria, ParAB*S* system (Badrinarayanan *et al.*, 2015, Wang *et al.*, 2013) and the Structural Maintenance of Chromosomes (SMC) condensin-like complex (Badrinarayanan *et al.*, 2015, Dame *et al.*, 2020, Nolivos & Sherratt, 2014).

ParAB*S* systems are widespread in bacteria (Livny *et al.*, 2007) and consist of three components: Firstly, one or more *parS* sequences located close to *ori* (Livny *et al.*, 2007). Secondly, the ParB protein, which binds *parS* sequences and then spreads along the chromosome to form large nucleoprotein complexes (Lin & Grossman, 1998, Mohl & Gober, 1997, Graham *et al.*, 2014, Murray *et al.*, 2006). Thirdly, the P-loop ATPase ParA, which binds DNA non-specifically and also interacts with the ParB·*parS* complex (Leonard *et al.*, 2005, Ptacin *et al.*, 2010, Fogel & Waldor, 2006, Schofield *et al.*, 2010, Lim *et al.*, 2014). ParA binds DNA in its ATP-bound dimeric form, and ParB stimulates the low intrinsic ATPase activity of ParA leading to the formation of ParA monomers, which do not bind DNA (Leonard *et al.*, 2005, Ptacin *et al.*, 2010, Schofield *et al.*, 2010, Lim *et al.*, 2014). The mechanism of ParABS systems in chromosome segregation is best understood in *C. crescentus* (Ptacin *et al.*, 2010, Schofield *et al.*, 2010, Lim *et al.*, 2014). Here, upon replication and formation of the two *ori*-proximal ParB·*parS* complexes, one remains at the pole and the other interacts with ParA bound non-specifically to the chromosome. This interaction results in stimulation of ParA ATPase activity and its release from the chromosome, leaving ParB·*parS* complex free to interact with adjacent DNA-bound ParA dimers. Successive cycles of these interactions generates a gradient of ParA across the nucleoid and results in ParB·*parS* complex translocation across the nucleoid towards the opposite pole (Ptacin *et al.*, 2010, Schofield *et al.*, 2010, Lim *et al.*, 2014, Hwang *et al.*, 2013, Vecchiarelli *et al.*, 2013). Some ParAB*S* systems function together with polar landmark proteins to anchor ParB·*parS* complexes at the poles and sequester ParA (Bowman *et al.*, 2008, Ebersbach *et al.*, 2008, Schofield *et al.*, 2010, Ptacin *et al.*, 2014, Yamaichi *et al.*, 2012, Donovan *et al.*, 2012, Lin *et al.*, 2017). Interestingly, these landmark proteins are much less conserved than the ParAB proteins. Generally, a ParAB*S* system is dispensable for viability but lack thereof causes chromosome segregation defects resulting in the formation of anucleate cells (Vallet-Gely & Boccard, 2013, Fogel & Waldor, 2006, Donovan *et al.*, 2010, Ginda *et al.*, 2013, Yamaichi *et al.*, 2007, Lee & Grossman, 2006) and only in a few species has this system been shown to be essential (Mohl & Gober, 1997, Harms *et al.*, 2013, Iniesta, 2014, Jung *et al.*, 2019).

In bacteria, three types of SMC complexes with low sequence similarity have been identified, i.e. the Smc/ScpA/ScpB, MukB/MukE/MukF and MksB/MksE/MksF complexes (Nolivos & Sherratt, 2014) with the Smc/ScpA/ScpB type being the most widespread (Gruber, 2011). Smc proteins have an ATPase head formed by the N-and C-terminal domains, a hinge, and a long anti-parallel coiled-coil domain that separates ATPase head and hinge (Nolivos & Sherratt, 2014, Yatskevich *et al.*, 2019) (Fig. 1A). The hinge mediates dimerization and ATP binding results in formation of a ring-shaped Smc dimer (Nolivos & Sherratt, 2014, Yatskevich *et al.*, 2019). The ScpA kleisin homolog reinforces ring formation while the ScpB dimer interacts with ScpA (Nolivos & Sherratt, 2014, Yatskevich *et al.*, 2019) (Fig. 1A). Based on work in several bacteria, SMC complexes of the Smc/ScpA/ScpB type are loaded onto the two daughter chromosomes during chromosome segregation at the ParB·*parS* complexes close to *ori* (Minnen *et al.*, 2011, Sullivan *et al.*, 2009, Gruber & Errington, 2009, Tran *et al.*, 2017, Böhm *et al.*, 2020). Each SMC complex is thought to encircle the two replichores of a daughter chromosome and translocate many kbp away from the loading site while aligning the two replichores and, in parallel, individualizing the two daughter chromosomes (Minnen *et al.*, 2016, Tran *et al.*, 2017, Le *et al.*, 2013, Wang *et al.*, 2017, Wang *et al.*, 2015, Marbouty *et al.*, 2015, Böhm *et al.*, 2020). Inactivation of SMC typically causes conditional growth defects associated with chromosome segregation defects whereby cells are viable at slow growth rates but have growth defects or even suffer lethality at increased growth rates (Britton *et al.*, 1998, Le *et al.*, 2013, Gruber *et al.*, 2014, Wang *et al.*, 2014, Minnen *et al.*, 2011, Mascarenhas *et al.*, 2002, Soppa *et al.*, 2002, Moriya *et al.*, 1998, Yu *et al.*, 2010).

**Figure 1.**
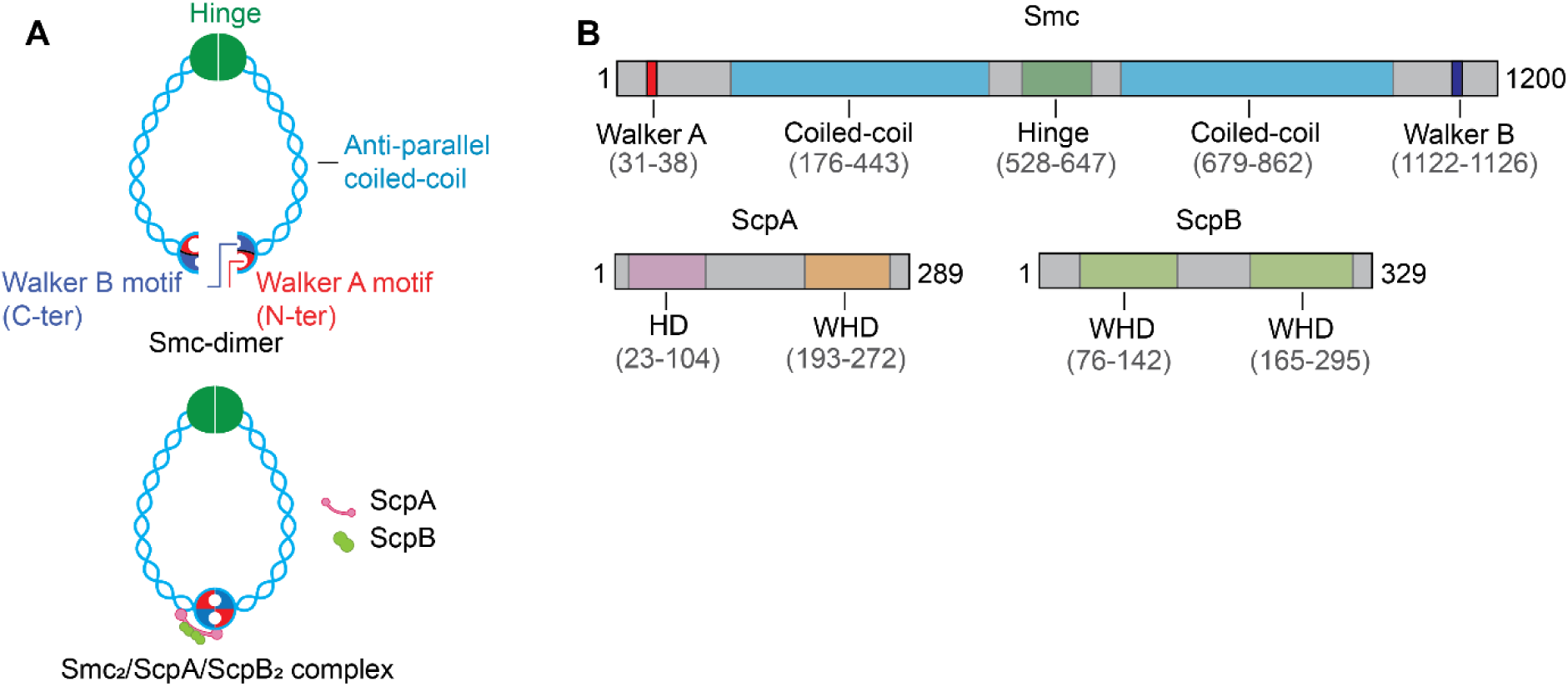
Domain architecture of proteins of the SMC complex in *M. xanthus.* (A) Architecture of the Smc dimer and Smc_2_ScpAScpB_2_ complex. (B) Domain architecture of *M. xanthus* Smc, ScpA and ScpB. In Smc, the five conserved domains or sequence motifs. ScpA has the characteristic domain organization with an N-terminal helical domain (HD) and a C-terminal winged-helix domain (WHD) (Fig. S2A) while ScpB contains the two characteristic WHD domains (Fig. S2B) (Yatskevich *et al.*, 2019).

*Myxococcus xanthus* is a Gram-negative, rod-shaped deltaproteobacterium with a unique lifecycle in which transitions between different stages are regulated by nutrient availability (Konovalova *et al.*, 2010). In the presence of nutrients, newborn cells contain a single 9.14 Mb sized chromosome (Goldman *et al.*, 2006) that is replicated once per cell cycle (Harms *et al.*, 2013, Iniesta, 2014) and spores, which are formed in response to starvation, contain two densely packed chromosomes (Tzeng & Singer, 2005). Newborn cells have an *ori-ter* longitudinal arrangement in which *ori* and *ter* are oriented towards the old and new pole, respectively. Uniquely, *ori* and *ter* localize subpolarly and are separated from the tip of the poles by ∼1 µm (Harms *et al.*, 2013, Iniesta, 2014). Specifically, the ParB·*parS* complexes localize ∼1 µm from the pole tips while ParA forms large subpolar patches bridging from a pole tip to a subpolar ParB·*parS* complex (Harms *et al.*, 2013, Iniesta, 2014). These distinct patterns depend on the three bactofilins BacN, BacO, BacP and the PadC adaptor protein (Lin *et al.*, 2017). Bactofilins polymerize spontaneously *in vitro* (Kühn *et al.*, 2010, Bulyha *et al.*, 2013, Koch *et al.*, 2011, Vasa *et al.*, 2015, Deng *et al.*, 2019) and PadC is a ParB-like protein (Lin *et al.*, 2017, Osorio-Valeriano *et al.*, 2019). *In vivo* and *in vitro* evidence supports that BacNOP together with PadC co-assemble to form subpolar scaffolds extending ∼1 µm away from a pole tip to anchor the ParB *parS* complexes at their pole-distal end, thereby, robustly positioning these complexes at a distance from the pole tip. Monomeric ParA interacts with PadC in the BacNOP/PadC scaffold and localizes along the entire scaffold (Lin *et al.*, 2017, Osorio-Valeriano *et al.*, 2019). ParA and ParB are essential, and their depletion causes chromosome segregation defects and divisions over the nucleoid (Iniesta, 2014, Harms *et al.*, 2013). BacNOP and PadC are not essential but in their absence the ParB·*parS* complexes localize more randomly, ParA is no longer sequestered in the subpolar regions, the nucleoid is reduced in length, and cells have a moderate increase in chromosome number (Lin *et al.*, 2017).

To understand how the unique chromosome arrangement in *M. xanthus* is accomplished, we focused on the function of SMC in chromosome segregation and organization. We show that SMC is important for both processes and is conditionally essential with mutants lacking SMC being non-viable at an increased temperature. We also report that BacNOP/PadC scaffold is important for chromosome segregation. Finally, lack of both SMC and the BacNOP/PadC scaffold is synthetic lethal altogether supporting a model whereby these two systems have distinct and partially redundant functions in chromosome segregation and organization *M. xanthus.*

## Results

### Deletion of *smc* or *scpAB* causes a temperature sensitive growth defect

We identified *M. xanthus* homologs of Smc (MXAN_4901), ScpA (MXAN_3841) and ScpB (MXAN_3840) but not of MukBEF or MksBEF types of SMC complexes (Fig. 1B; Fig. S1-2). Orthologs of *M. xanthus* Smc, ScpA and ScpB with high sequence identity/similarity were identified in Myxoccocales with fully sequenced genomes (Fig. S3AB) and the neighborhood of the *M. xanthus* genes was conserved in other Myxococcales (Fig. S3AB). The genetic organization suggests that *smc* is not part of an operon while the 3’ end of *scpA* and 5’ end of *scpB* overlap indicating co-transcription (Fig. S3AB).

To assess the function of Smc, ScpA and ScpB in *M. xanthus* we considered that *smc* mutations can cause temperature sensitive growth defects being lethal at increased temperatures. We attempted to generate an in-frame deletion of *smc* and a double *scpAB* in-frame deletion in the wild-type (WT) DK1622 at 25°C as well as at 32°C, the latter being the temperature at which *M. xanthus* is normally propagated under laboratory conditions. Both mutants were readily obtained at 25°C but not at 32°C. By contrast, in strains containing a plasmid integrated in a single copy at the *attB* site and expressing either *smc-mCherry* or *scpAB* form the native promoter (Fig. S3AB), we readily obtained the in-frame deletions at 32°C. WT and the two complementation strains had similar growth rates in casitone growth medium at 25°C while the two in-frame deletion mutants had a reduced growth rate; at 32°C, growth of the Δ*smc* and Δ*scpAB* mutants dramatically decreased immediately after the temperature shift (Fig. 2A). The temperature sensitivity of the Δ*smc* and Δ*scpAB* mutants was also observed on solid casitone medium (Fig. 2A). Immunoblots showed that the Smc-mCherry fusion protein accumulated as a full-length protein in the complementation strain (Fig. S4A) documenting that the fusion protein is active.

**Figure 2.**
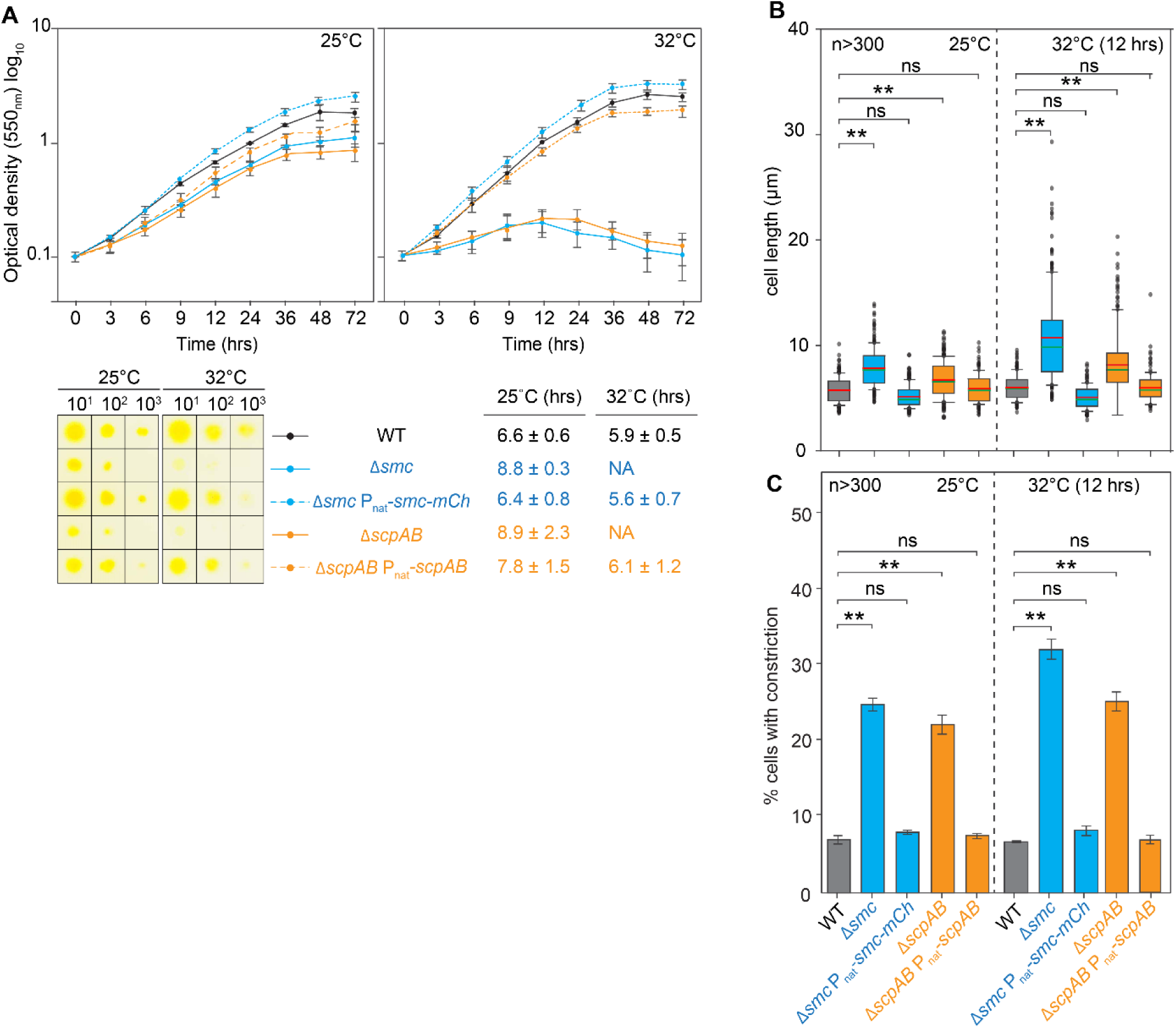
Growth and cell division defects in Δ*smc* and Δ*scpAB* cells. (A) Growth curves of indicated strains at 25°C and 32°C in suspension (upper) and on a solid surface (lower). In the growth curves, mean ± standard deviation (SD) are shown from three biological replicates. In the plating assay, plates were incubated for 48 hrs. Table indicates generation times of strains at indicated temperatures, NA, not applicable. (B) Cell length distribution of cells of indicated genotypes at 25°C and 32°C (12 hrs). In the box plots, red and green indicate the median and mean, respectively, boxes the 25^th^ and 75^th^ percentiles, and whiskers the 10^th^ and 90^th^ percentile. n>300 for all strains from three biological replicates. ** p <0.001, ns, not significant in Mann-Whitney test. (C) Quantitation for cell division constrictions in cells of indicated genotypes at 25°C and 32°C (12 hrs). Same cells as in (B) were analyzed. ** p <0.001, ns, not significant in one-sided Student’s t-test.

Phase contrast microscopy revealed that Δ*smc* and Δ*scpAB* cells at 25°C were significantly longer than WT cells, and after 12 hrs at 32°C (from hereon 32°C (12 hrs)), Δ*smc* and Δ*scpAB* cells were even longer (Fig. 2B; Fig. S5). At both temperatures, an increased fraction of Δ*smc* and Δ*scpAB* cells had a cell division constriction at midcell compared to WT but the mutants did not form minicells (Fig. 2C; Fig. S5) suggesting that completion of cell division was delayed in both mutants while positioning of the division site was unaffected. The cell length and division defects were complemented by ectopic expression of *smc-mCherry* and *scpAB* in the relevant strains (Fig. 2BC; Fig. S5). We conclude that Smc and ScpAB are conditionally essential and that lack of Smc or ScpAB causes a temperature sensitive growth defect with lethality at 32°C that is associated with a cell division defect.

### Lack of SMC causes aberrant chromosome segregation

To examine the role of SMC in chromosome segregation and organization, we determined the DNA content of cells by flow cytometry. At 25°C and 32°C, WT and the two complementation strains contained one-to-two chromosomes in agreement with previous observations (Tzeng & Singer, 2005). Δ*smc* and Δ*scpAB* cells at 25°C had a decreased fraction of cells with one chromosome and at 32°C, almost all cells contained two chromosomes and some also had an abnormally high DNA content (Fig. 3A). Consistently, DAPI (4′,6-diamidino-2-phenylindole)-stained cells of WT and the two complementation strains contained one or two nucleoids at 25°C and 32°C (Fig. 3B). In agreement with previous reports (Harms *et al.*, 2013, Treuner-Lange *et al.*, 2013, Schumacher *et al.*, 2017), single nucleoids were centered around midcell while in cells with two segregated nucleoids, these were centered around the quarter cell length positions (Fig. 3B) and in these cells constrictions were visible between segregated nucleoids (Fig. 3B). By contrast, many Δ*smc* and Δ*scpAB* cells at 25°C had a single nucleoid mass centered around midcell; at 32° (12 hrs), the nucleoids were also not well separated and more randomly localized but mostly centered at midcell (Fig. 3B). No significant differences in nucleoid length were observed between WT and Δ*smc* and Δ*scpAB* cells at 25°C and 32°C (Fig. 3C). In agreement with these observations, Δ*smc* and Δ*scpAB* cells often had large subpolar regions devoid of DNA (Fig. 3B). At both temperatures, cell division constrictions in Δ*smc* and Δ*scpAB* cells were frequently observed over unsegregated nucleoids (Fig. 3B). Because Δ*smc* and Δ*scpAB* cells at both temperatures are enriched for cells with two chromosomes (Fig. 3A), these observations support that Smc and ScpAB are important for chromosome segregation but not for nucleoid condensation. Because Δ*smc* and Δ*scpAB* cells are viable at 25°C, we infer that the chromosomes eventually segregate; the delayed chromosome segregation would explain why these cells are longer and have a higher frequency of cells with cell division constrictions. At 32°C, we infer that the segregation defect is more dramatic and, therefore, cells do not divide.

**Figure 3.**
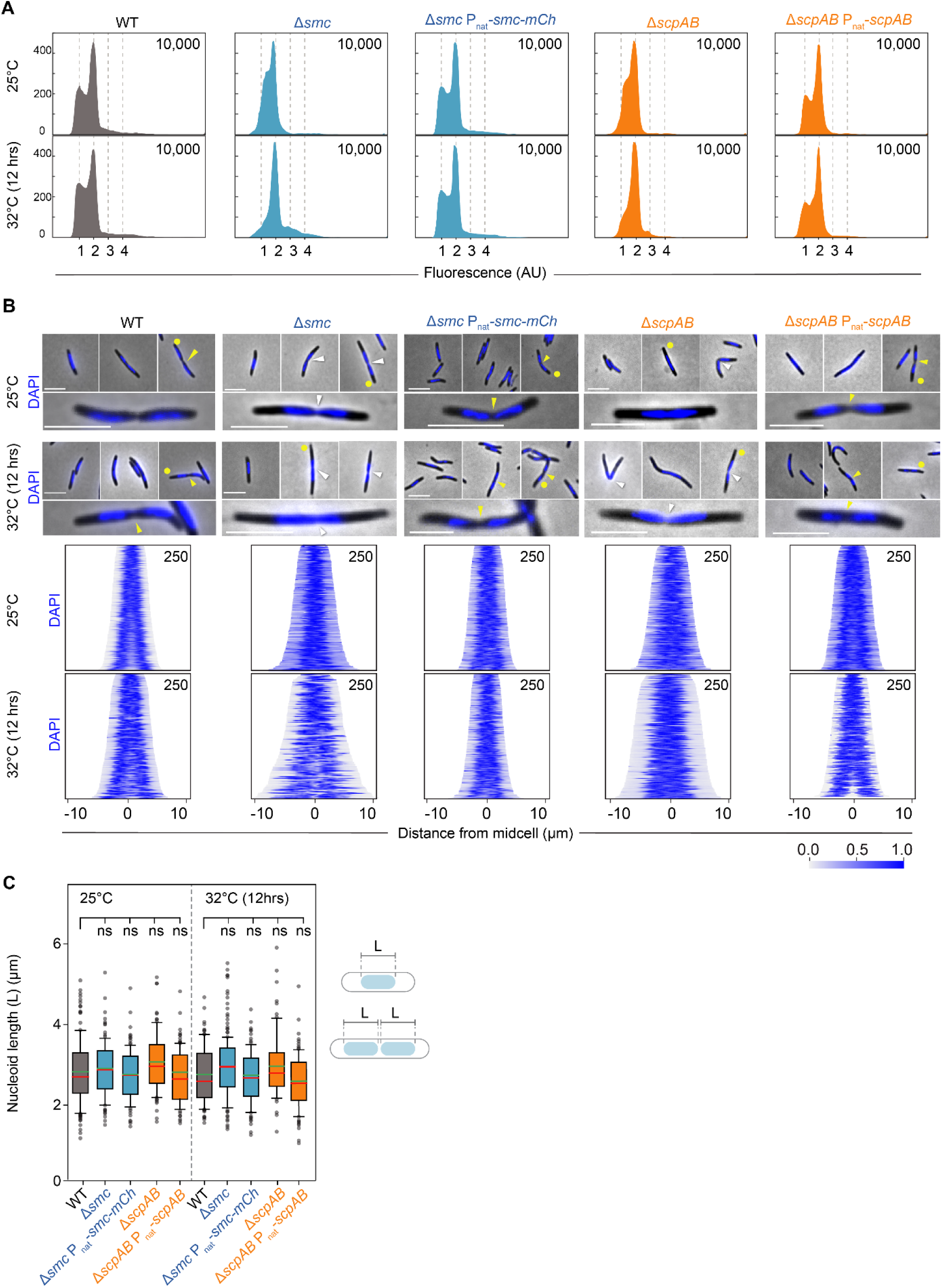
Smc and ScpAB are important for chromosome segregation. (A) Flow-cytometry analysis for DNA content in cells of indicated genotypes. Cells were stained with Vybrant® DyeCycle™ Orange and 10,000 cells analysed per strain. Y-axis, number of cells and X-axis, fluorescence in arbitrary units (AU). Numbers below X-axis indicate chromosome numbers. (B) Nucleoid staining of cells of the indicated genotypes. Cells grown at 25°C and 32°C (12 hrs) were stained with DAPI (blue). Upper panels, yellow dots indicate magnified cells; yellow and white arrowheads indicate constrictions between segregated nucleoids and over nucleoids, respectively. Scale bars, 5 µm. Lower panels, demographs of DAPI stained cells at 25°C and 32°C (12 hrs). Cell profiles were sorted according to cell length and in random orientation. Numbers in upper right, number of cells included from three biological replicates. (C) Analysis of nucleoid length in cells of indicated genotypes. DAPI stained cells from (B) were analyzed for nucleoid length. Only nucleoids that were not visibly undergoing segregation were included (see schematics on right). Box plot as in Fig. 2B; ns, not significant in Mann-Whitney test.

### Smc and ScpAB are important for positioning of ParB·*parS* complexes and *ter*

To further investigate chromosome segregation and organization in cells lacking SMC, we visualized *ori* using the ParB *parS* complex and *ter* using a fluorescence reporter operator system (FROS)-marker with a *tetO* array at 180° (from hereon FROS-180°) as proxies (Harms *et al.*, 2013). ParB binds to 24 *parS* sites located ∼30 kb from *ori* (Treuner-Lange *et al.*, 2013, Harms *et al.*, 2013, Iniesta, 2014) forming a large nucleoprotein complex that covers ∼20 kb (Skotnicka *et al.*, 2020). ParB-YFP and TetR-YFP, which binds the *tet*O array, were synthesized from plasmids integrated at the *attB* site and accumulated at 25°C and 32°C in all strains tested (Fig. S6AB).

At both temperatures, WT and the complementation strains contained one or two ParB-YFP foci in the subpolar regions and at the “edge” of nucleoids (Fig. 4AB; Fig. S7A). At 25°C, Δ*smc* and Δ*scpAB* cells, generally, had one or two ParB-YFP foci in the subpolar regions and only a few cells with >2 foci. By contrast, at 32°C (12 hrs), and in agreement with an increase in the chromosome content as shown by flow cytometry, a significantly higher number of cells had >2 ParB-YFP foci and, generally, the foci were more randomly distributed than at 25°C (Fig. 4AB; Fig. S7A). At both temperatures, the shortest distance from a ParB-YFP focus to the pole tip was significantly increased compared to WT (Fig. S7B) in agreement with the observations that Δ*smc* and Δ*scpAB* cells often had large subpolar regions devoid of DNA (Fig. 3B).

**Figure 4.**
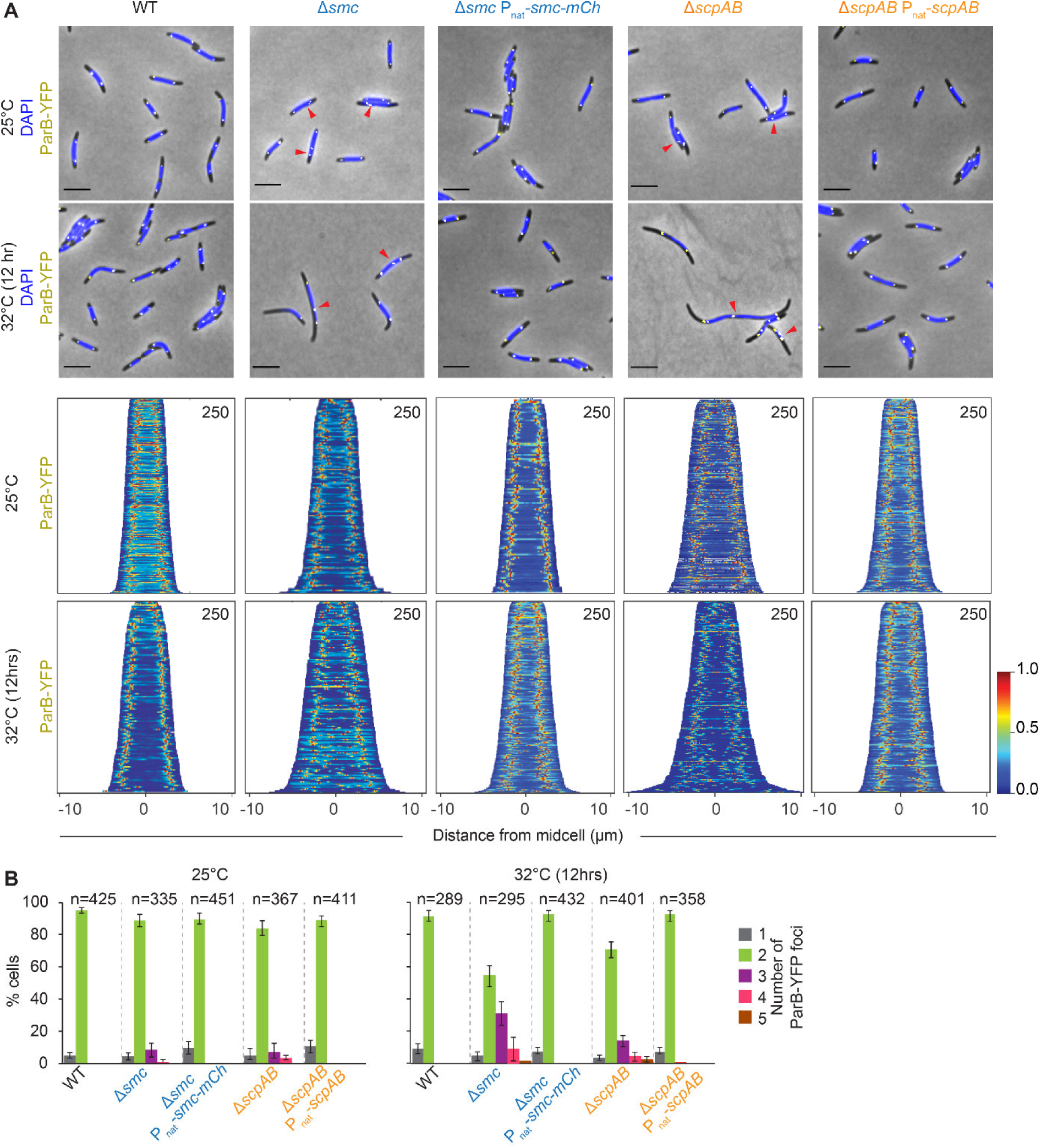
Smc and ScpAB are important for positioning of ParB·*parS* complexes. (A) ParB-YFP localization in cells of indicated genotypes. ParB-YFP (yellow) was expressed ectopically from the native *parB* promoter in merodiploid *parB*^+^/*parB-YFP* strains. Cells were DAPI-stained (blue) and visualized by phase contrast and epifluorescence microscopy and merged images generated. Red arrowsheads, cells with >2 ParB-YFP foci. Scale bars, 5 µm. Lower panels: Demographs of ParB-YFP signals. Cell profiles were sorted according to cell size and in random cell orientation. Numbers, number of cells included from three biological replicates. (B) Quantitation of ParB foci. Cells were treated as in (A) and the number of ParB-YFP foci per cell quantified. N, number of cells analysed from three biological replicates. Error bars, SD.

At both temperatures, the large majority of WT cells had one FROS-180° focus somewhere over the nucleoid and <5% of cells had two foci, which were located in both cell halves around midcell (Fig. 5AB; Fig. S7C) as previously described (Harms *et al.*, 2013). By comparison, significantly more Δ*smc* and Δ*scpAB* cells at both temperatures had two foci (Fig. 5AB) and, these foci were more randomly organized than in WT and frequently not in opposite cell halves (Fig. 5A; Fig. S7C).

**Figure 5.**
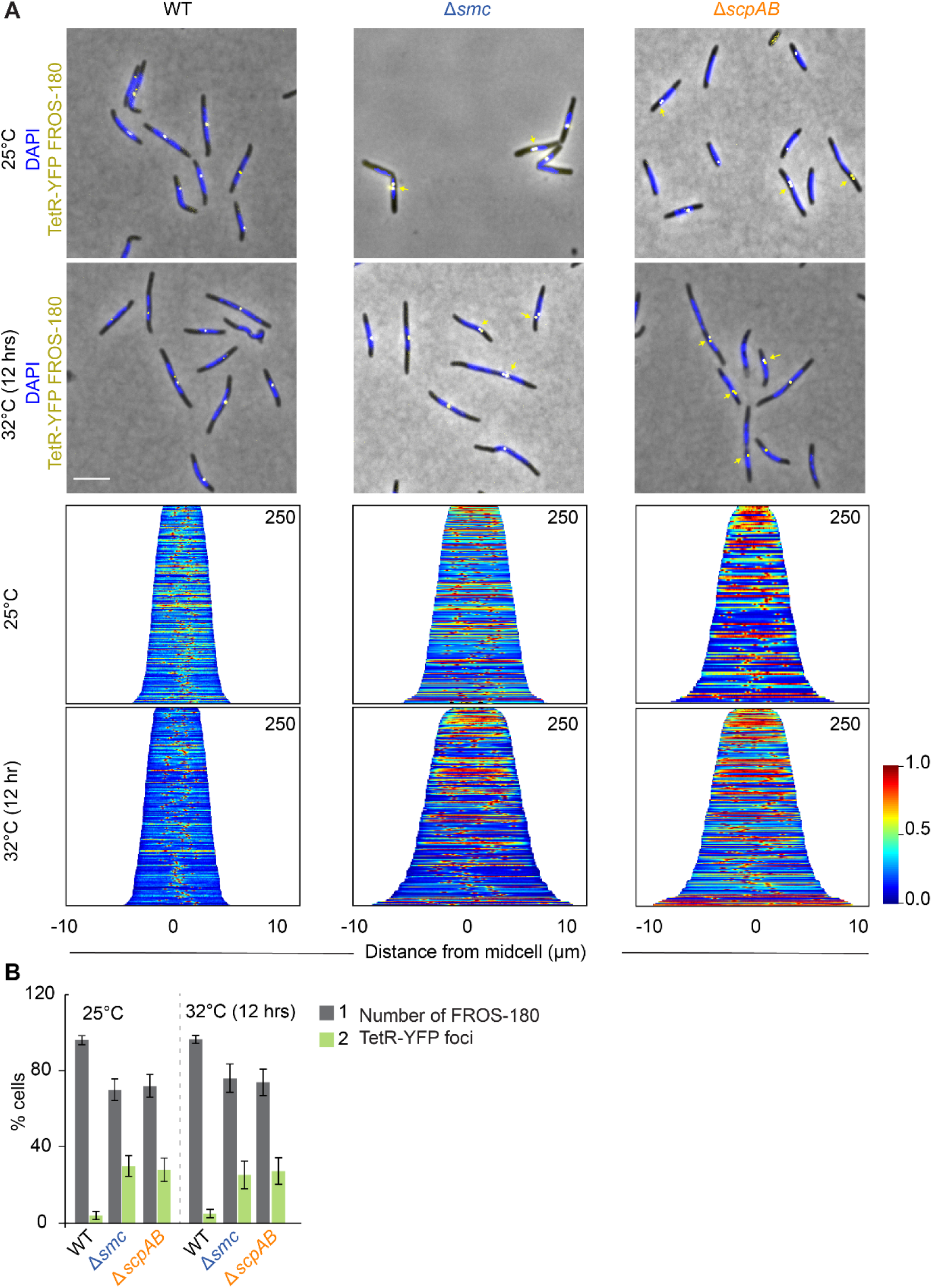
Smc and ScpAB are important for positioning of *ter*. (A) FROS-180° was visualized by binding of TetR-YFP in cells of indicated genotypes. *tetR-YFP* was expressed from the *cuo* promoter. Cells were grown in the presence of 300 µM CuSO_4_ for 6 hrs before microscopy to induce synthesis of TetR-YFP (yellow). Nucleoids were stained with DAPI (blue) and cells imaged by phase contrast and epifluorescence microscopy. Yellow arrows cells with mislocalized FROS-180° focus. Scale bars, 5 µm. Lower panels, demographs of TetR-YFP signal. Cell profiles were sorted according to cell size and in random cell orientation. Numbers in upper right, number of cells included from three biological replicates. (B) Quantification of TetR-YFP foci. Cells from (A) were analysed for the number of TetR-YFP foci per cell and plotted according to the colour code on the right. Error bars, SD.

We conclude that SMC is important for positioning of ParB·*parS* complexes and *ter* consistent with a defect in chromosome segregation. Moreover, we speculate that the increased fraction of Δ*smc* cells with two *ter* regions is a consequence of the segregation defect and, consequently, a delay in cell division.

### Lack of SMC causes a defect in ParB·*parS* complex segregation

To specifically address a segregation defect caused by lack of SMC, we performed time-lapse fluorescence microscopy at 25°C and 32°C on WT and the Δ*smc* mutant containing ParB-YFP. Because ParB·*parS* complexes segregate early during the cell cycle (Harms *et al.*, 2013), we focused on cells that underwent division and then followed ParB-YFP in these cells. In WT at 25°C/32°C, duplication of the ParB-YFP focus was evident at 25±8/20±4 min (n=12/15) after cell division and while one focus remained in the subpolar region of the old pole, the second translocated directionally to the opposite subpolar region in 27±9/26±8 min. Both foci remained stably localized in the subpolar regions after segregation (Fig. 6A).

**Figure 6.**
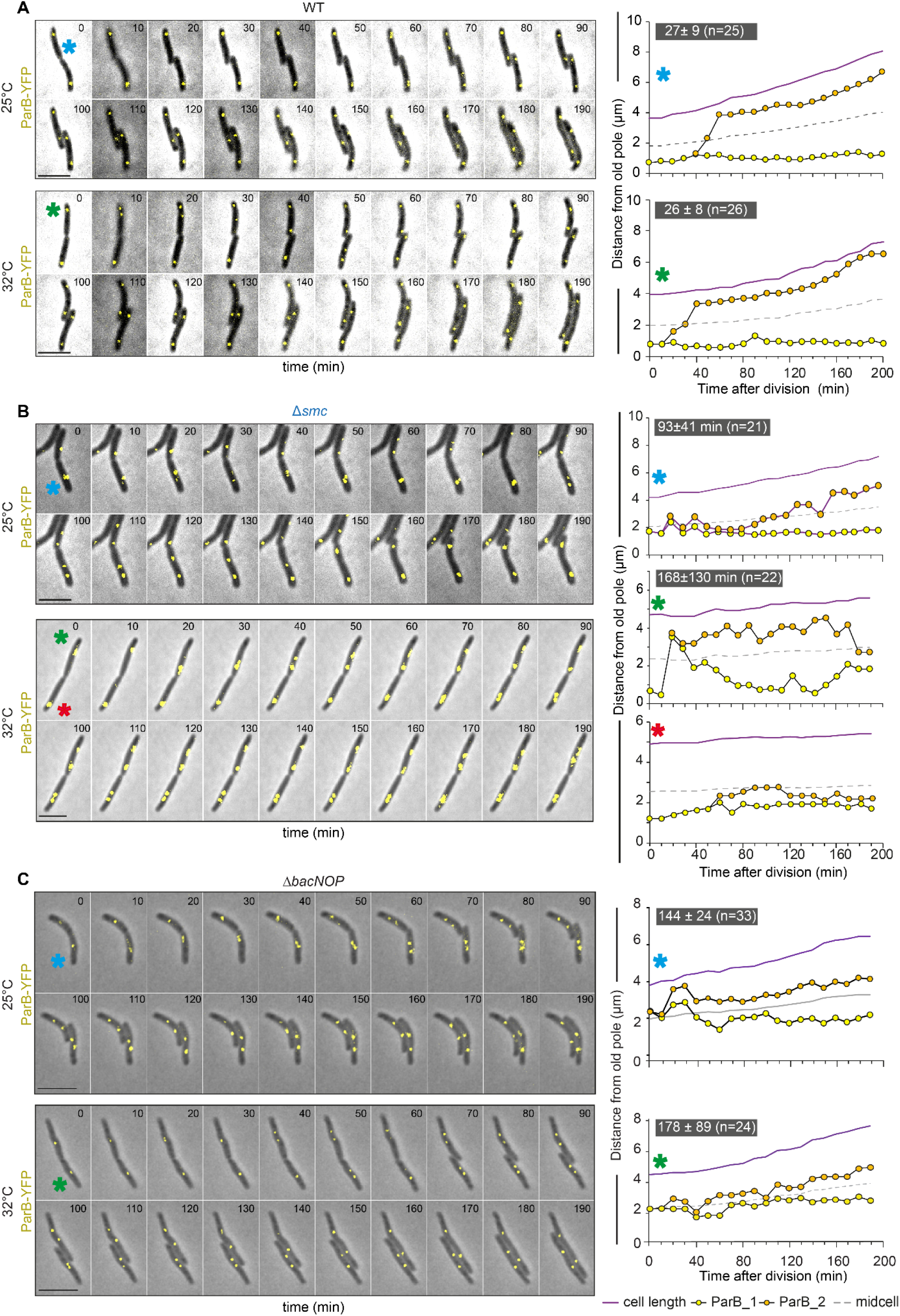
Lack of Smc or BacNOP causes a ParB·*parS* complex segregation defect. (A, B, C) Cells were grown at 25°C and transferred to a 1.0% agarose pad supplemented with 0.2% casitone and imaged after 15 min and then every 10 min at 25°C and 32°C. Montages (left panels) and line graphs of ParB-YFP trajectories (yellow, orange) (right panels) are shown. ParB-YFP was expressed ectopically from the native *parB* promoter in merodiploid *parB*^+^/*parB-YFP* strains. Colored asterisks in montages indicate the cell for which ParB-YFP trajectories are shown on the right. Scale bars, 5 µm. Average time ± SD for translocation of ParB-YFP focus from the old to the subpolar region at the new pole is indicated in white on the grey background together with the number of cells analysed. All strains contained the Δ*mglA* mutation to inactivate the motility systems.

In Δ*smc* cells at 25°C, duplication of the ParB-YFP focus was evident at 25±12 min (n=16) after cell division and the translocating cluster moved aberrantly with frequent pauses and reversals reaching the opposite subpolar region in 93±41 min (Fig. 6B). At 32°C, cluster duplication was evident after 68±27 min (n=24) and the translocating cluster moved to the opposite subpolar region occurred in 168±130 min and with frequent pauses and reversals (Fig. 6B); in 8% of cells, none of the clusters translocated across midcell (Fig. 6B, cell marked with red asterisk). In addition, we frequently observed that ParB·*parS* foci were not stably localized in the subpolar regions (Fig. 6B, cell marked with green asterisk) in agreement with the more random localization of these foci by snap-shot analysis (Fig. 4A; S7A). Because the flow cytometry data (Fig. 3A) and snap-shot analysis of ParB·*parS* complexes (Fig. 4AB; S7AB) and the *ter* region (Fig. 5AB; S7C) do not support that lack of SMC causes a replication initiation defect, we speculate that the longer time interval between cell division and evident cluster duplication is caused by the segregation defect rather than a defect in initiation of replication.

We conclude that SMC is important for ParB·*parS* complex segregation and that lack of SMC causes delayed, aberrant and sometimes failed ParB·*parS* complex segregation and a defect in anchoring of ParB·*parS* complexes in the subpolar regions.

### Subpolar ParA-mCherry, PadC-YFP and BacP localization and organization are disturbed in the absence of Smc and ScpAB

Because ParB·*parS* segregation depends on ParA and positioning at a distance from the pole tips depends on the BacNOP/PadC scaffold, we examined the accumulation and localization of ParA, PadC and BacP in the absence of Smc or ScpAB.

As reported (Harms *et al.*, 2013, Iniesta, 2014, Lin *et al.*, 2017), in WT cells at both temperatures, ParA-mCherry not only displayed the asymmetric signal over the nucleoid indicative of a segregating ParB·*parS* complex but also the characteristic patches in the nucleoid-free subpolar regions; and, while smaller cells had unipolar localization of ParA-mCherry, the remaining cells had bipolar localization (Fig. 7A). By contrast, in Δ*smc* and Δ*scpAB* cells at 25°C as well as at 32°C, the subpolar ParA-mCherry patches were of lower intensity and appeared fragmented (Fig. 7A). ParA and ParA-mCherry accumulated at the same level in all three strains at both temperatures (Fig. S8A).

**Figure 7.**
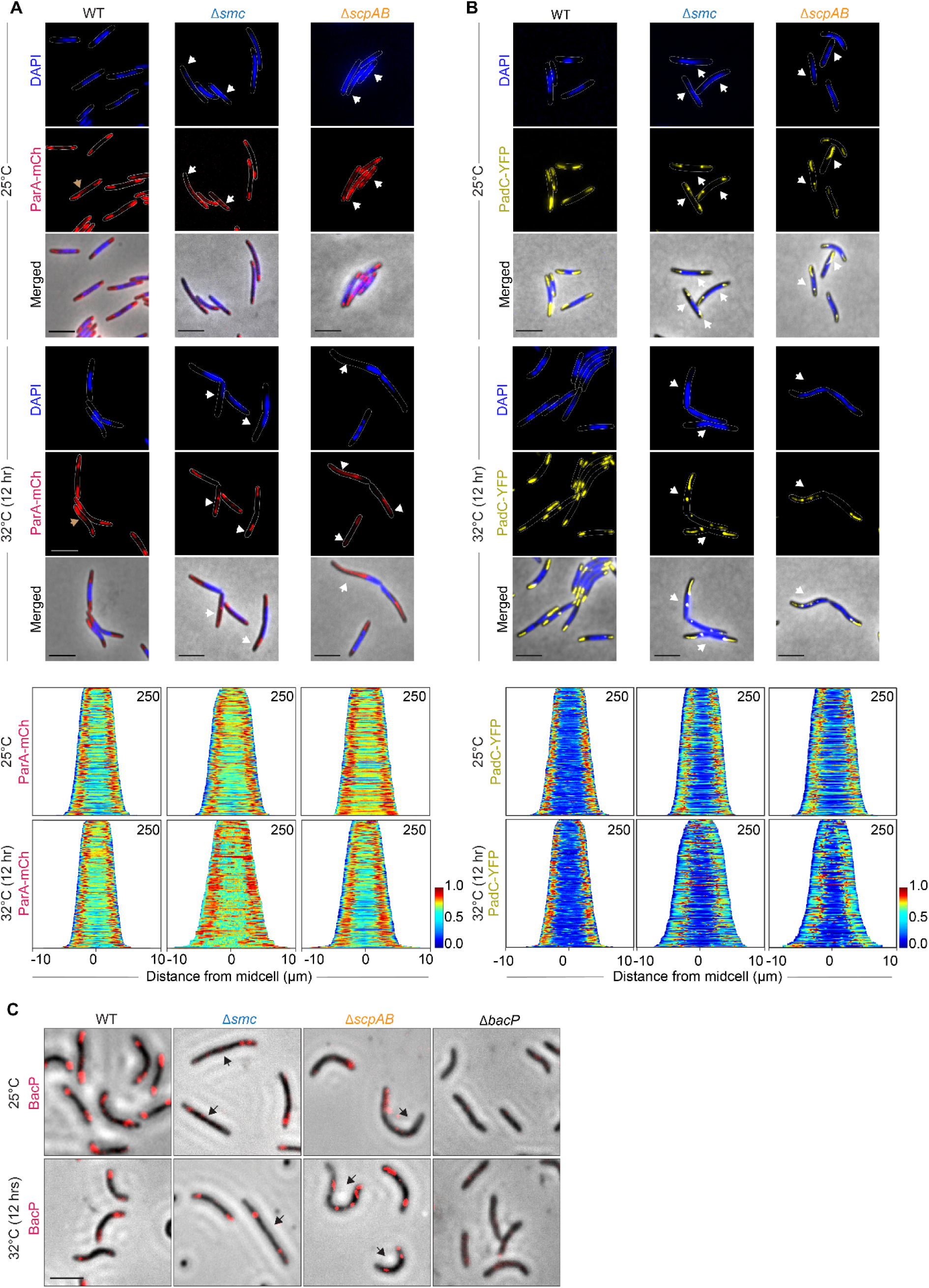
Smc and ScpAB are important for subpolar localization of ParA-mCherry, PadC-YFP and BacP. (A, B) WT, Δ*smc* and Δ*scpAB* cells expressing ParA-mCherry from the native promoter at the *attB* site in *parA*^+^/*parA-mCherry* merodiploids (A) or PadC-YFP from a vanillate inducible promoter in *padC*^*+*^*/padC-YFP* merodiploids (B) were grown at 25°C and 32°C (12 hrs) and imaged. Top row, DAPI stain (blue), middle row ParA-mCherry (red) and PadC-YFP (yellow), lower row merged images. In (A), brown arrows indicate asymmetric cloud of ParA-mCherry over the nucleoid; white arrows indicate cells with abnormal ParA-mCherry or PadC-YFP localization. Scale bars, 5 µm. Lower panels, demographs of ParA-mCherry and PadC-YFP. Numbers, number of cells included from three biological replicates. (C) Immuno-fluorescence microscopy of BacP in WT, Δ*smc*, Δ*scpAB* and Δ*bacP* cells. The Δ*bacP* mutant was used as negative control. Black arrows indicate cells with abnormal bacP localilzation. Scale bar, 5µm.

As previously reported (Lin *et al.*, 2017), PadC-YFP in WT occupied both subpolar regions at 25°C and 32°C and was occasionally also detected in a cluster over the nucleoid (Fig. 7B). By contrast, in Δ*smc* and Δ*scpAB* cells at 25°C and 32°C the patches were of lower intensity and appeared fragmented and, generally, with strong signals at the edge of nucleoids and sometimes over nucleoids. PadC and PadC-YFP accumulated similarly in all three strains at both temperatures (Fig. S8B).

To examine BacNOP, we focused on BacP, which is the only of the three BacNOP bactofilins that is absolutely essential for assembly of the BacNOP/PadC complex (Lin *et al.*, 2017), and determined its localization using immuno-fluorescence microscopy. BacP accumulated similarly in all three strains at both temperatures (Fig. S8C). In WT at 25°C and at 32°C, BacP localized to the subpolar regions similar to ParA and PadC in agreement with previous observations (Bulyha *et al.*, 2013, Lin *et al.*, 2017). However, in Δ*smc* and Δ*scpAB* cells, the BacP signal in the subpolar regions was also fragmented at both temperatures (Fig. 7C).

We conclude that SMC is important for localization and organization of ParA-mCherry, PadC-YFP and BacP in the subpolar regions at 25°C and 32°C. We speculate that this effect causes the reduced anchoring of the ParB·*parS* complexes in the subpolar regions in the absence of SMC.

### Lack of the BacNOP/PadC scaffold causes a defect in ParB·*parS* complex segregation

Next, we analyzed whether the BacNOP/PadC scaffold is also important for chromosome segregation. At 25°C and 32°C, Δ*bacNOP* and Δ*padC* cells had the same time interval from division to visible duplication of the ParB·*parS* focus as WT (Δ*bacNOP* at 25°/32°C: 26±6/27±10 min (n=20/25); Δ*padC* at 25°/32°C: 26±9/23±7 min (n=22/23)). Both mutants displayed the longer distance between ParB·*parS* foci and the tip of the poles at both temperatures as described (Lin *et al.*, 2017). More importantly, the two mutants had ParB·*parS* complex segregation defects with ParB·*parS* complex translocation times of ranging from 144±24 to 201±61 min (Fig. 6C; S9). Because Smc-mCherry accumulated at WT levels in the Δ*bacNOP* and Δ*padC* cells (Fig. S4C) these observations support that the BacNOP/PadC scaffold is not only important for the subpolar anchoring of ParB·*parS* complexes and subpolar ParA localization but also for chromosome segregation.

### Deletion of *smc/scpAB* in combination with *bacNOP* or *padC* is synthetic lethal

Lack of the BacNOP/PadC scaffold affects chromosome segregation, anchoring of ParB·*parS* complexes and subpolar ParA localization but is not essential for viability (Lin *et al.*, 2017). As shown here, lack of Smc or ScpAB is conditional lethal causing chromosome segregation and organization defects and interfering with the subpolar localization/organization of BacP, PadC and ParA. To test whether the BacNOP/PadC scaffold and SMC function redundantly in chromosome segregation and organization, we tested specific double mutants for synthetic lethality at 25° (Table 1).

**Table 1.** Inactivation of SMC is synthetic lethal with lack of the BacNOP/PadC scaffold

| Strain genotype | Target gene(s) | Deletion of target gene(s) obtained |
| --- | --- | --- |
| WT | *bacNOP* | Yes |
| $\Delta smc$ | *bacNOP* | No |
| $\Delta smc, P_{nat-smc-mCh}$ | *bacNOP* | Yes |
| $\Delta scpAB$ | *bacNOP* | No |
| DK1622 | *padC* | Yes |
| $\Delta smc$ | *padC* | No |
| $\Delta smc, P_{nat-smc-mCh}$ | *padC* | Yes |
| $\Delta scpAB$ | *padC* | No |
| $\Delta bacNOP$ | *smc* | No |

*bacNOP* could easily be deleted *en bloc* in WT; however, in the Δ*smc* and Δ*scpAB* mutants, we only obtained the *bacNOP* mutation in the presence of complementing *smc-mCherry* or *scpAB* constructs, respectively expressed ectopically. Similar results were obtained for deletion of *padC*. Finally, we observed that *smc* could not be deleted in the Δ*bacNOP* mutant. Thus, simultaneous loss of SMC and BacNOP/PadC cause synthetic lethality indicating that the two systems function redundantly to support chromosome organization and segregation.

### Smc forms multiple dynamic foci over the nucleoid

Smc has been reported to form dynamic foci over the nucleoid in several bacteria (Minnen *et al.*, 2011, Jensen & Shapiro, 2003, Mascarenhas *et al.*, 2002). Therefore, we analyzed the localization of Smc in *M. xanthus* using the active Smc-mCherry fusion. Snap-shot images demonstrated that Smc-mCherry formed 1 to 6 foci per cell over the nucleoid in the Δ*smc* strain as well as in the *smc*^+^/*smc-mCherry* merodiploid (Fig. 8AB). The number of foci correlated slightly with the cell length (Fig. S10A) but the foci did not localize in a recognizable pattern (Fig. S10B). Smc-mCherry cluster formation was not observed in the Δ*scpAB* mutant (Fig. S10C) despite Smc-mCherry accumulating in this strain (Fig. S4B) supporting that the foci represent assembled and active SMC complexes.

**Figure 8.**
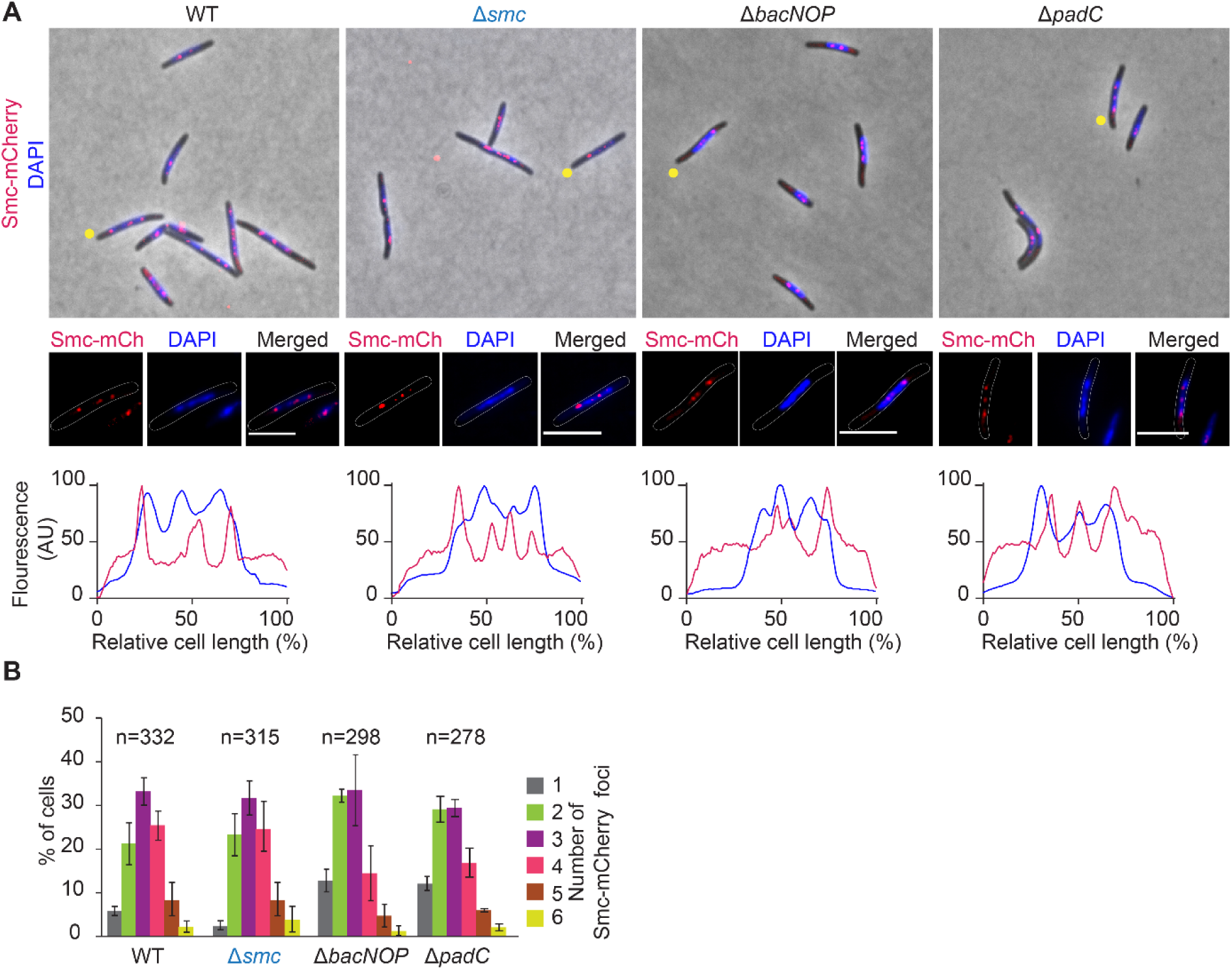
Smc-mCherry forms foci over the nucleoid. (A) Localization of Smc-mCherry in strains of indicated genotypes. All strains except for the Δ*smc* strain, are *smc*^+^/*smc-mCherry* merodiploids and all strains express *smc-mCherry* ectopically from the native promoter at the *attB* site. Cells were grown at 32°C. Upper panels, cells were stained with DAPI (blue) and imaged. Shown are merged images of phase contrast, DAPI and Smc-mCherry signals. Below, representative cells (marked with yellow dot in upper panels) with DAPI and Smc-mCherry signals separately and merged; below, line scans of the representative cells. Scale bars, 5 µm. (B) Analysis of number of Smc-mCherry foci per cell. In cells from A, the number of Smc-mCherry foci per cell was quantified. N, number of cells analysed from three biological replicates. Error bars, SD.

Smc-mCherry accumulated in the Δ*padC* and Δ*bacNOP* mutants and formed foci over the nucleoid as in WT (Fig. 8AB; Fig. S4C; Fig. S10A). We conclude that Smc-mCherry forms foci independently of BacNOP and PadC.

Time-lapse fluorescence microscopy of the Δ*smc smc-mCherry* strain demonstrated that the Smc-mCherry foci were relocating dynamically, sometimes giving the impression that they would split; however, overall, no recognizable pattern was evident (Fig. S10D). We conclude that Smc forms dynamic clusters over the nucleoid.

### Smc-mCherry is unstable in the absence of ParB

Because ParB·*parS* complexes are important for loading of SMC, we investigated the relationship between ParB and SMC in *M. xanthus*. First, we asked whether the two proteins colocalize. To this end, Smc-mCherry and ParB-YFP were co-expressed in the Δ*smc* mutant. Accumulation of both fusions proteins was confirmed by immunoblot analysis (Fig. S11A). Epifluorescence microscopy showed that both proteins formed foci of which only a few overlapped (Fig. S11B). Consistently, time-lapse microscopy demonstrated that Smc-mCherry formed dynamic foci independently of where ParB-YFP foci localized (Fig. S11C).

To analyze the localization of Smc-mCherry in the absence of ParB, Smc-mCherry was expressed from its native promoter from the ectopic *MXAN_18-19* site while *parB* was expressed from the Cu^2+^ inducible *cuo* promoter from the *attB* site in a Δ*parB* mutant. In the presence of 300 µM CuSO_4_, both proteins were detected in immunoblots (Fig. S12). Upon removal of CuSO_4_, ParB was depleted, and at 18 hrs largely no longer detected. Interestingly, Smc-mCherry was no longer detectable at 9 hrs of ParB depletion suggesting that ParB is important for Smc stability. Because Smc-mCherry was no longer detectable at the earliest timepoint where ParB was largely depleted, we did not determine Smc-mCherry localization in the absence of ParB.

## Discussion

Here, we identify the SMC condensin-like complex composed of Smc and ScpAB in *M. xanthus* and demonstrate that it is conditionally essential. Mutants lacking either the Smc subunit or the ScpAB subunits of SMC were viable at 25°C but not at 32°C. At the permissive temperature, cells had chromosome segregation defects, were slightly longer than WT, and had an increased frequency of cells with cell division constrictions. At the non-permissive temperature, these defects were exacerbated. Because cells in which cell division is inhibited are filamentous but do not have chromosome segregation defects (Schumacher *et al.*, 2017, Treuner-Lange *et al.*, 2013), we conclude that the primary defect in the Δ*smc* and Δ*scpAB* mutants is in chromosome segregation and organization and not in cell division. Consistently, direct measurement of the segregation kinetics of ParB·*parS* complexes demonstrated that they segregate more slowly and aberrantly in the absence of the SMC complex than in WT. Moreover, our result support that the Δ*smc* and Δ*scpAB* mutants are non-viable at 32°C because cells cannot divide due to the unsegregated chromosomes.

*M. xanthus* cells use the ParAB*S* for chromosome segregation and the BacNOP/PadC scaffold to anchor the ParB·*parS* complexes and ParA at distinct locations in the subpolar regions (Lin *et al.*, 2017, Harms *et al.*, 2013, Iniesta, 2014). By analysing ParB·*parS* complex segregation kinetics, we found that the BacNOP/PadC scaffold is not only involved in positioning of ParAB*S* components but also important for chromosome segregation. Thus, three systems are important for chromosome segregation in *M. xanthus*, i.e. the ParAB*S* system, the BacNOP/PadC scaffold and SMC. Interestingly, lack of any single one of the three causes distinct phenotypes. ParA and ParB are essential and lack of ParA or ParB causes chromosome segregation defects, cell divisions over the nucleoid and the formation of chromosome free cells (Harms *et al.*, 2013, Iniesta, 2014). By contrast, the BacNOP bactofilins and PadC are not essential and lack of BacNOP or PadC causes a slight increase in chromosome content, segregation defects, loss of the subpolar anchoring of the ParB·*parS* segregation complexes, lack of ParA localization in the subpolar regions, and shorter nucleoids [here, (Lin *et al.*, 2017, Osorio-Valeriano *et al.*, 2019)]. Finally, lack of SMC is conditionally lethal and causes chromosome segregation and organization defects as well as inhibition of cell division. We observed that Δ*smc* Δ*bacNOP*, Δ*scpAB* Δ*bacNOP*, Δ*smc* Δ*padC* and Δ*scpAB* Δ*padC* double mutants were synthetic lethal. These observations together with the different phenotypes of mutants lacking SMC or BacNOP/PadC support that these two systems function redundantly to support chromosome segregation and organization, yet with distinct functions in these processes. Because cells that lack SMC are viable at 25°C but not at 32°C while cells lacking SMC as well as the BacNOP/PadC scaffold are not viable under any condition tested, we conclude that the BacNOP/PadC scaffold is functional at 25°C in the absence of SMC but not at 32°. Along the same lines, SMC is functional at both temperatures in the absence of the BacNOP/PadC scaffold. Thus, SMC can still exert its essential function in the absence of the BacNOP/PadC system.

What, then, is the function of these three systems and how are they connected? Because the ParAB*S* system is essential, we suggest that this system makes up the basic machinery for chromosome segregation in *M. xanthus*. BacNOP and PadC form a subpolar scaffold in a mutually dependent manner, i.e. lack BacNOP results in mislocalization of PadC and *vice versa* (Lin *et al.*, 2017). The BacNOP/PadC scaffold is important for the subpolar positioning of the ParB·*parS* segregation complexes and sequestration of monomeric forms of ParA *via* binding to PadC (Lin *et al.*, 2017). As shown here, BacNOP/PadC is not important for SMC accumulation and localization suggesting that this scaffold is also not important for chromosomal loading of SMC. We speculate that the segregation defect in the absence of the BacNOP/PadC scaffold is caused by lack of ParA sequestration, which would interfere with formation of the ParA gradient across the nucleoid that is important for directional ParB·*parS* segregation (Ptacin *et al.*, 2010, Schofield *et al.*, 2010). In this scenario, the BacNOP/PadC scaffold functions to make the ParABS system operate more robustly analogously to the polar landmark proteins PopZ and TipN in *C. crescentus*, which anchors ParB·*parS* segregation complexes (Bowman *et al.*, 2008, Ebersbach *et al.*, 2008) and sequesters ParA (Schofield *et al.*, 2010, Ptacin *et al.*, 2014), respectively. Like in several other bacteria, our data are consistent with the notion that the primary function of SMC is in directly individualizing the two daughter chromosomes during chromosome segregation. Lack of SMC also caused aberrant localization of BacP, PadC and ParA, which might also contribute to the chromosome segregation defect. How this effect is brought about is not known but could be a consequence of the increased length of the nucleoid-free subpolar regions in the absence of SMC or, alternatively, a direct interaction between BacNOP/PadC and SMC. In several bacteria, the ParB·*parS* complex serves as a loading platform for SMC (Gruber & Errington, 2009, Minnen *et al.*, 2011, Sullivan *et al.*, 2009, Tran *et al.*, 2017). We observed that depletion of ParB caused instability of Smc supporting that the two proteins may interact; however, it remains to be determined whether ParB is important for loading of SMC in *M. xanthus*.

Altogether, we suggest a model whereby the large 9.14 Mb *M. xanthus* chromosome is segregated and laid down in a spatially organized manner by three interating systems, ParAB*S* constitutes the basic machinery for chromosome segregation while SMC and BacNOP/PadC have distinct yet redundant roles in this process with SMC supporting individualization of daughter chromosomes and BacNOP/PadC making the ParABS system operate more robustly.

## Materials and methods

### *M. xanthus* and *E.coli* strains and growth

*M. xanthus* strains used in this study are all derivatives of the WT DK1622 (Kaiser, 1979). *M. xanthus* strains, plasmids and oligonucleotides used are listed in Table 2, 3 and S1, respectively. *M. xanthus* cells were grown in liquid 1% CTT medium or on 1% CTT/1.5% agar plates at 32°C (Hodgkin & Kaiser, 1977). Kanamycin and oxytetracycline were added to *M. xanthus* cells at concentrations of 40 µg/ml or 10 µg/ml, respectively. In-frame gene deletions were generated as described previously (Shi *et al.*, 2008). All strains were tested by PCR. Vanillate and CuSO_4_ were added where indicated at the indicated concentrations. Growth was measured as an increase in OD at 550 nm. *E. coli* strains were grown in LB broth in the presence of relevant antibiotics (Sambrook & Russell, 2001). All plasmids were propagated in *E. coli* Mach1 (Δ*recA*1398 *endA*1 *tonA* Φ80Δ*lacM*15 Δ*lacX*74 *hsdR*(r_K_^-^ m_K_^+^)).

**Table 2.**
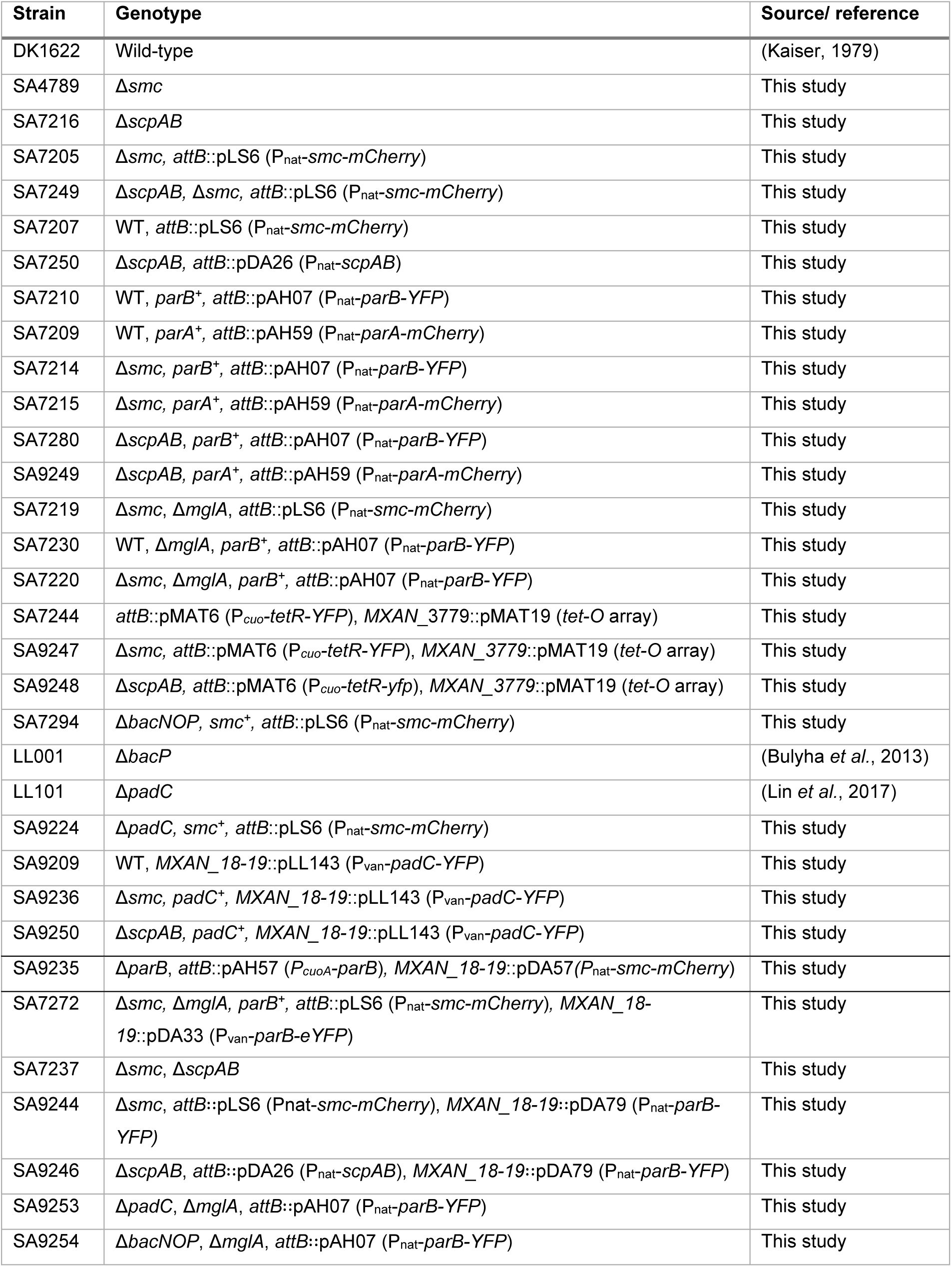
*M. xanthus* strains used in this study

**Table 3.** List of plasmids used in this study

| Plasmid | Feature | Reference |
| --- | --- | --- |
| pBJ114 | Plasmid for generation of in-frame deletions in *M. xanthus*; *kanR*<sup>+</sup>, *galK*<sup>+</sup>, | (Julien *et al.*, 2000) |
| pSWU30 | Plasmid for integration at Mx8 *attB* site, Tet | (Wu *et al.*, 1997) |
| pMR3690 | Plasmid for integration at *MXAN_18-19* site; vanillate inducible promoter for gene expression | (Iniesta *et al.*, 2012) |
| pAH07 | Derivative of pSWU30, P<sub>nat</sub>-*parB*-YFP | (Harms *et al.*, 2013) |
| pAH59 | Derivative of pSWU30, P<sub>nat</sub>-*parA*-mCherry | (Harms *et al.*, 2013) |
| pDS27 | pBJ114; construct for in-frame deletion of *smc* (*MXAN_4901*) | This study |
| pLS6 | Derivative of pSWU30, P<sub>nat</sub>-*smc*-mCherry | This study |
| pDA11 | pBJ114; construct for in-frame deletion of Δ*scpAB* (*MXAN_3841*, *MXAN_3840*) | This study |
| pMAT19 | pBJ114; construct for integration of *tetO* array at *MXAN_3779* | (Harms *et al.*, 2013) |
| pMAT6 | P<sub>cuo</sub>-*tetR*-YFP | (Harms *et al.*, 2013) |
| pSL16 | pBJ114, Δ*mglA* | (Miertzschke *et al.*, 2011) |
| pDA26 | pSWU30, P<sub>nat</sub>-*scpAB* | This study |
| pDA33 | pMR3690, P<sub>van</sub>-*parB*-eYFP | This study |
| pMT982 | Construct for in-frame deletion of *bacNOP* (*MXAN_4637-4635*) | (Kühn *et al.*, 2010) |
| pLL38 | Construct for in-frame deletion of *padC* (*MXAN_4634*) | (Lin *et al.*, 2017) |

### Fluorescence microscopy and live cell imaging

Exponentially growing cells were transferred to slides with a thin pad of 1.0% agarose (SeaKem LE agarose, Cambrex) with TPM buffer (10 mM Tris-HCl pH 7.6, 1 mM KH_2_PO_4_ pH 7.6, 8 mM MgSO_4_), covered with a coverslip and imaged with a temperature-controlled Leica DMi8 inverted microscope. Phase contrast and fluorescence images were acquired using a Hamamatsu ORCA-flash V2 Digital CMOS camera. Time-lapse imaging was performed as described (Schumacher & Søgaard-Andersen, 2018). Briefly, cells were transferred to pre-warmed 1% agarose pads supplemented with 0.2% casitone in TPM buffer mounted on a metallic microscopy slide. Slides were covered with parafilm to retain humidity of the agarose. Cells in phase contrast images were automatically detected using Oufti (Paintdakhi *et al.*, 2016) or the MicrobeJ plugin (Ducret *et al.*, 2016) in ImageJ (Schneider *et al.*, 2012). Kymographs were generated using MicrobeJ and ImageJ. Cell length and fluorescence signals were extracted using one of the above-mentioned applications. For DAPI staining of nucleoids, cells were incubated with 0.5 µg/ml DAPI for 10 min prior to microscopy. Image processing was performed using Metamorph v7.5 (Molecular Devices). Immunofluorescence microscopy detection of BacP was performed as described (Bulyha *et al.*, 2013).

### Flow cytometry

Exponentially growing cells (OD_550_ 0.5-0.7) were stained using Vybrant® DyeCycle™ Orange Stain (VDCO) (Invitrogen, cat number V35005) at a final concentration of 50 µM at 25°C and 32°C for 30 min according to the sample treatment. Cells were diluted 10-fold in TPM buffer before DNA content measurement by flow cytometry (BD LSR Fortessa, Becton Dickinson GmbH, Heidelberg, Germany). 10,000 cells were analysed for each data set. Three technical and three biological triplicates were analysed per strain.

### Immunoblot analysis

Immunoblots were carried out as described (Sambrook & Russell, 2001). α-BacP (Bulyha *et al.*, 2013), α-PilC (Bulyha *et al.*, 2009), α-ParB (Harms *et al.*, 2013), α-ParA (Harms *et al.*, 2013), α-PadC (Lin *et al.*, 2017), α-mCherry (Biovision), and α-GFP (Roche Diagnostics GmbH, Mannheim, Germany) antibodies were used together with horseradish-conjugated goat α-rabbit (Sigma-Aldrich,Germany) or horseradish-conjugated α-mouse sheep IgG antibody as secondary antibody. Blots were developed using Luminata Crescendo Western HRP Substrate (Millipore) and visualized using a LAS-4000 luminescent image analyzer (Fujifilm).

### Bioinformatics

*M. xanthus* Smc, ScpA and ScpB were identified using KEGG orthology (Kanehisa & Goto, 2000) and BLASTP (Boratyn *et al.*, 2013) searches with Smc, ScpA and ScpB from *C. crescentus* and *B. subtilis* as queries. Protein domains were assigned using SMART (Simple Modular Architecture Research Tool) (Letunic & Bork, 2018), NCBI conserved domain (Marchler-Bauer *et al.*, 2017) and KEGG (Kanehisa & Goto, 2000). Protein sequences were aligned with Clustal X2.1 (Larkin *et al.*, 2007) and modified using Genedoc (Nicholas *et al.*, 1997).

### Statistics

Statistical analyses were performed using SigmaPlot v14. Data sets were tested for normal distribution using a Shapiro-Wilk test. Student’s *t*-test was used for data with a normal distribution and a Mann-Whitney test in the case of a non-uniform distribution of data.

## Supporting information

All supplementary data

## Acknowledgements

We thank Dobromir Szadkowski and Dorota Skotnicka for helpful discussions and Lara Stühn for help with construction of plasmids and strains. This work was supported by the Deutsche Forschungsgemeinschaft (DFG) within the framework of the Transregio 174 “Spatiotemporal dynamics of bacterial cells” and the Max Planck Society.

## Data availability

The data that support the findings of this study are available on request from the corresponding author.

## References

Badrinarayanan, A., Le, T.B.K., and Laub, M.T. (2015) Bacterial chromosome organization and segregation. Ann. Rev. Cell Dev. Biol. 31: 171–199.

Böhm, K., Giacomelli, G., Schmidt, A., Imhof, A., Koszul, R., Marbouty, M., and Bramkamp, M. (2020) Chromosome organization by a conserved condensin-ParB system in the actinobacterium *Corynebacterium glutamicum*. Nat. Comm. 11: 1485.

Boratyn, G.M., Camacho, C., Cooper, P.S., Coulouris, G., Fong, A., Ma, N., Madden, T.L., Matten, W.T., McGinnis, S.D., Merezhuk, Y., Raytselis, Y., Sayers, E.W., Tao, T., Ye, J., and Zaretskaya, I. (2013) BLAST: a more efficient report with usability improvements. Nucleic Acids Res. 41: W29–33.

Bowman, G.R., Comolli, L.R., Zhu, J., Eckart, M., Koenig, M., Downing, K.H., Moerner, W.E., Earnest, T., and Shapiro, L. (2008) A polymeric protein anchors the chromosomal origin/ParB complex at a bacterial cell pole. Cell 134: 945–955.

Britton, R.A., Lin, D.C., and Grossman, A.D. (1998) Characterization of a prokaryotic SMC protein involved in chromosome partitioning. Genes Dev 12: 1254–1259.

Bulyha, I., Lindow, S., Lin, L., Bolte, K., Wuichet, K., Kahnt, J., van der Does, C., Thanbichler, M., and Søgaard-Andersen, L. (2013) Two small GTPases act in concert with the bactofilin cytoskeleton to regulate dynamic bacterial cell polarity. Dev Cell 25: 119–131.

Bulyha, I., Schmidt, C., Lenz, P., Jakovljevic, V., Hone, A., Maier, B., Hoppert, M., and Søgaard-Andersen, L. (2009) Regulation of the type IV pili molecular machine by dynamic localization of two motor proteins. Mol Microbiol 74: 691–706.

Dame, R.T., Rashid, F.-Z.M., and Grainger, D.C. (2020) Chromosome organization in bacteria: mechanistic insights into genome structure and function. Nat. Rev. Genet. 21: 227–242.

Deng, X., Gonzalez Llamazares, A., Wagstaff, J.M., Hale, V.L., Cannone, G., McLaughlin, S.H., Kureisaite-Ciziene, D., and Löwe, J. (2019) The structure of bactofilin filaments reveals their mode of membrane binding and lack of polarity. Nat. Microbiol. 4: 2357–2368.

Donovan, C., Schwaiger, A., Krämer, R., and Bramkamp, M. (2010) Subcellular localization and characterization of the ParAB System from *Corynebacterium glutamicum*. J. Bacteriol. 192: 3441–3451.

Donovan, C., Sieger, B., Kramer, R., and Bramkamp, M. (2012) A synthetic *Escherichia coli* system identifies a conserved origin tethering factor in Actinobacteria. Mol Microbiol 84: 105–116.

Ducret, A., Quardokus, E.M., and Brun, Y.V. (2016) MicrobeJ, a tool for high throughput bacterial cell detection and quantitative analysis. Nat Microbiol 1: 16077.

Ebersbach, G., Briegel, A., Jensen, G.J., and Jacobs-Wagner, C. (2008) A self-associating protein critical for chromosome attachment, division, and polar organization in *Caulobacter*. Cell 134: 956–968.

Fogel, M.A., and Waldor, M.K. (2006) A dynamic, mitotic-like mechanism for bacterial chromosome segregation. Genes Dev. 20: 3269–3282.

Ginda, K., Bezulska, M., Ziólkiewicz, M., Dziadek, J., Zakrzewska-Czerwińska, J., and Jakimowicz, D. (2013) ParA of *Mycobacterium smegmatis* co-ordinates chromosome segregation with the cell cycle and interacts with the polar growth determinant DivIVA. Mol. Microbiol. 87: 998–1012.

Goldman, B.S., Nierman, W.C., Kaiser, D., Slater, S.C., Durkin, A.S., Eisen, J.A., Ronning, C.M., Barbazuk, W.B., Blanchard, M., Field, C., Halling, C., Hinkle, G., Iartchuk, O., Kim, H.S., Mackenzie, C., Madupu, R., Miller, N., Shvartsbeyn, A., Sullivan, S.A., Vaudin, M., Wiegand, R., and Kaplan, H.B. (2006) Evolution of sensory complexity recorded in a myxobacterial genome. Proc Natl Acad Sci U S A 103: 15200–15205.

Graham, T.G., Wang, X., Song, D., Etson, C.M., van Oijen, A.M., Rudner, D.Z., and Loparo, J.J. (2014) ParB spreading requires DNA bridging. Genes Dev 28: 1228–1238.

Gruber, S. (2011) MukBEF on the march: taking over chromosome organization in bacteria? Mol. Microbiol. 81: 855–859.

Gruber, S., and Errington, J. (2009) Recruitment of condensin to replication origin regions by ParB/SpoOJ promotes chromosome segregation in *B. subtilis*. Cell 137: 685–696.

Gruber, S., Veening, J.-W., Bach, J., Blettinger, M., Bramkamp, M., and Errington, J. (2014) Interlinked sister chromosomes arise in the absence of condensin during fast replication in *B. subtilis*. Curr. Biol. 24: 293–298.

Harms, A., Treuner-Lange, A., Schumacher, D., and Søgaard-Andersen, L. (2013) Tracking of chromosome and replisome dynamics in *Myxococcus xanthus* reveals a novel chromosome arrangement. PLoS Genet 9: e1003802.

Hodgkin, J., and Kaiser, D. (1977) Cell-to-cell stimulation of movement in nonmotile mutants of *Myxococcus*. Proc Natl Acad Sci USA 74: 2938–2942.

Hwang, L.C., Vecchiarelli, A.G., Han, Y.W., Mizuuchi, M., Harada, Y., Funnell, B.E., and Mizuuchi, K. (2013) ParA-mediated plasmid partition driven by protein pattern self-organization. EMBO J 32: 1238–1249.

Iniesta, A.A. (2014) ParABS system in chromosome partitioning in the bacterium *Myxococcus xanthus*. PLoS One 9: e86897.

Iniesta, A.A., García-Heras, F., Abellón-Ruiz, J., Gallego-García, A., and Elías-Arnanz, M. (2012) Two systems for conditional gene expression in *Myxococcus xanthus* inducible by isopropyl-β-thiogalactopyranoside or vanillate. J. Bacteriol. 194: 5875–5885.

Jensen, R.B., and Shapiro, L. (2003) Cell-cycle-regulated expression and subcellular localization of the *Caulobacter crescentus* SMC chromosome structural protein. J Bacteriol 185: 3068–3075.

Julien, B., Kaiser, A.D., and Garza, A. (2000) Spatial control of cell differentiation in Myxococcus xanthus. Proc Natl Acad Sci U S A 97: 9098–9103.

Jung, A., Raßbach, A., Pulpetta, R.L., van Teeseling, M.C.F., Heinrich, K., Sobetzko, P., Serrania, J., Becker, A., and Thanbichler, M. (2019) Two-step chromosome segregation in the stalked budding bacterium *Hyphomonas neptunium*. Nat. Comm. 10: 3290.

Kaiser, D. (1979) Social gliding is correlated with the presence of pili in *Myxococcus xanthus*. Proc Natl Acad Sci USA 76: 5952–5956.

Kanehisa, M., and Goto, S. (2000) KEGG: kyoto encyclopedia of genes and genomes. Nucleic Acids Res 28: 27–30.

Koch, M.K., McHugh, C.A., and Hoiczyk, E. (2011) BacM, an N-terminally processed bactofilin of *Myxococcus xanthus*, is crucial for proper cell shape. Mol. Microbiol. 80: 1031–1051.

Konovalova, A., Petters, T., and Søgaard-Andersen, L. (2010) Extracellular biology of *Myxococcus xanthus*. FEMS Microbiol. Rev. 34: 89–106.

Kühn, J., Briegel, A., Morschel, E., Kahnt, J., Leser, K., Wick, S., Jensen, G.J., and Thanbichler, M. (2010) Bactofilins, a ubiquitous class of cytoskeletal proteins mediating polar localization of a cell wall synthase in *Caulobacter crescentus*. EMBO J 29: 327–339.

Larkin, M.A., Blackshields, G., Brown, N.P., Chenna, R., McGettigan, P.A., McWilliam, H., Valentin, F., Wallace, I.M., Wilm, A., Lopez, R., Thompson, J.D., Gibson, T.J., and Higgins, D.G. (2007) Clustal W and Clustal X version 2.0. Bioinformatics 23: 2947–2948.

Le, T.B., Imakaev, M.V., Mirny, L.A., and Laub, M.T. (2013) High-resolution mapping of the spatial organization of a bacterial chromosome. Science 342: 731–734.

Lee, P.S., and Grossman, A.D. (2006) The chromosome partitioning proteins Soj (ParA) and Spo0J (ParB) contribute to accurate chromosome partitioning, separation of replicated sister origins, and regulation of replication initiation in *Bacillus subtilis*. Mol. Microbiol. 60: 853–869.

Leonard, T.A., Butler, P.J., and Löwe, J. (2005) Bacterial chromosome segregation: structure and DNA binding of the Soj dimer--a conserved biological switch. EMBO J 24: 270–282.

Letunic, I., and Bork, P. (2018) 20 years of the SMART protein domain annotation resource. Nucleic Acids Res 46: D493–D496.

Lim, H.C., Surovtsev, I.V., Beltran, B.G., Huang, F., Bewersdorf, J., and Jacobs-Wagner, C. (2014) Evidence for a DNA-relay mechanism in ParABS-mediated chromosome segregation. Elife 3: e02758.

Lin, D.C.-H., and Grossman, A.D. (1998) Identification and characterization of a bacterial chromosome partitioning site. Cell 92: 675–685.

Lin, L., Osorio Valeriano, M., Harms, A., Søgaard-Andersen, L., and Thanbichler, M. (2017) Bactofilin-mediated organization of the ParABS chromosome segregation system in *Myxococcus xanthus*. Nat Commun 8: 1817.

Livny, J., Yamaichi, Y., and Waldor, M.K. (2007) Distribution of centromere-like parS sites in bacteria: insights from comparative genomics. J Bacteriol 189: 8693–8703.

Marbouty, M., Le Gall, A., Cattoni, Diego I., Cournac, A., Koh, A., Fiche, J.-B., Mozziconacci, J., Murray, H., Koszul, R., and Nollmann, M. (2015) Condensin- and replication-mediated bacterial chromosome folding and origin condensation revealed by Hi-C and super-resolution maging. Mol Cell 59: 588–602.

Marchler-Bauer, A., Bo, Y., Han, L., He, J., Lanczycki, C.J., Lu, S., Chitsaz, F., Derbyshire, M.K., Geer, R.C., Gonzales, N.R., Gwadz, M., Hurwitz, D.I., Lu, F., Marchler, G.H., Song, J.S., Thanki, N., Wang, Z., Yamashita, R.A., Zhang, D., Zheng, C., Geer, L.Y., and Bryant, S.H. (2017) CDD/SPARCLE: functional classification of proteins via subfamily domain architectures. Nucleic Acids Res 45: D200–D203.

Mascarenhas, J., Soppa, J., Strunnikov, A.V., and Graumann, P.L. (2002) Cell cycle-dependent localization of two novel prokaryotic chromosome segregation and condensation proteins in *Bacillus subtilis* that interact with SMC protein. EMBO J 21: 3108–3118.

Miertzschke, M., Koerner, C., Vetter, I.R., Keilberg, D., Hot, E., Leonardy, S., Sogaard-Andersen, L., and Wittinghofer, A. (2011) Structural analysis of the Ras-like G protein MglA and its cognate GAP MglB and implications for bacterial polarity. EMBO J 30: 4185–4197.

Minnen, A., Attaiech, L., Thon, M., Gruber, S., and Veening, J.W. (2011) SMC is recruited to *oriC* by ParB and promotes chromosome segregation in *Streptococcus pneumoniae*. Mol Microbiol 81: 676–688.

Minnen, A., Burmann, F., Wilhelm, L., Anchimiuk, A., Diebold-Durand, M.L., and Gruber, S. (2016) Control of Smc Coiled Coil Architecture by the ATPase Heads Facilitates Targeting to Chromosomal ParB/parS and Release onto Flanking DNA. Cell Rep 14: 2003–2016.

Mohl, D.A., and Gober, J.W. (1997) Cell cycle–dependent polar localization of chromosome partitioning proteins in *Caulobacter crescentus*. Cell 88: 675–684.

Moriya, S., Tsujikawa, E., Hassan, A.K., Asai, K., Kodama, T., and Ogasawara, N. (1998) A Bacillus subtilis gene-encoding protein homologous to eukaryotic SMC motor protein is necessary for chromosome partition. Mol Microbiol 29: 179–187.

Murray, H., Ferreira, H., and Errington, J. (2006) The bacterial chromosome segregation protein Spo0J spreads along DNA from *parS* nucleation sites. Mol. Microbiol. 61: 1352–1361.

Nicholas, K.B., Nicholas Jr., H.B., and Deerfield II, D.W. (1997) Genedoc: Analysis and visualization of genetic variation. EMBNEW.NEWS 4: 14.

Nielsen, H.J., Ottesen, J.R., Youngren, B., Austin, S.J., and Hansen, F.G. (2006) The Escherichia coli chromosome is organized with the left and right chromosome arms in separate cell halves. Mol Microbiol 62: 331–338.

Niki, H., Yamaichi, Y., and Hiraga, S. (2000) Dynamic organization of chromosomal DNA in Escherichia coli. Genes Dev 14: 212–223.

Nolivos, S., and Sherratt, D. (2014) The bacterial chromosome: architecture and action of bacterial SMC and SMC-like complexes. FEMS Microbiol Rev 38: 380–392.

Osorio-Valeriano, M., Altegoer, F., Steinchen, W., Urban, S., Liu, Y., Bange, G., and Thanbichler, M. (2019) ParB-type DNA Segregation Proteins Are CTP-Dependent Molecular Switches. Cell 179: 1512-1524.e1515.

Paintdakhi, A., Parry, B., Campos, M., Irnov, I., Elf, J., Surovtsev, I., and Jacobs-Wagner, C. (2016) Oufti: an integrated software package for high-accuracy, high-throughput quantitative microscopy analysis. Mol Microbiol 99: 767–777.

Ptacin, J.L., Gahlmann, A., Bowman, G.R., Perez, A.M., von Diezmann, A.R.S., Eckart, M.R., Moerner, W.E., and Shapiro, L. (2014) Bacterial scaffold directs pole-specific centromere segregation. Proc. Natl. Acad. Sci. USA 111: E2046–E2055.

Ptacin, J.L., Lee, S.F., Garner, E.C., Toro, E., Eckart, M., Comolli, L.R., Moerner, W.E., and Shapiro, L. (2010) A spindle-like apparatus guides bacterial chromosome segregation. Nat Cell Biol 12: 791–798.

Sambrook, J.F., and Russell, D.W. (2001) Molecular Cloning - A Laboratory Manual. Cold Spring Harbor Laboratory Press, Cold Spring Harbor, New York.

Schneider, C.A., Rasband, W.S., and Eliceiri, K.W. (2012) NIH Image to ImageJ: 25 years of image analysis. Nat. Methods 9: 671–675.

Schofield, W.B., Lim, H.C., and Jacobs-Wagner, C. (2010) Cell cycle coordination and regulation of bacterial chromosome segregation dynamics by polarly localized proteins. EMBO J 29: 3068–3081.

Schumacher, D., Bergeler, S., Harms, A., Vonck, J., Huneke-Vogt, S., Frey, E., and Søgaard-Andersen, L. (2017) The PomXYZ proteins self-organize on the bacterial nucleoid to stimulate cell division. Dev Cell 41: 299–314 e213.

Schumacher, D., and Søgaard-Andersen, L. (2018) Fluorescence live-cell imaging of the complete vegetative cell cycle of the slow-growing social bacterium *Myxococcus xanthus*. J. Vis. Exp.: e57860.

Shi, X., Wegener-Feldbrugge, S., Huntley, S., Hamann, N., Hedderich, R., and Søgaard-Andersen, L. (2008) Bioinformatics and experimental analysis of proteins of two-component systems in *Myxococcus xanthus*. J Bacteriol 190: 613–624.

Skotnicka, D., Steinchen, D., Szadkowski, D., Cadby, I.T., Lovering, A.L., Bange, G., and Søgaard-Andersen, L. (2020) CdbA is a DNA-binding protein and c-di-GMP receptor important for nucleoid organization and segregation in *Myxococcus xanthus*. Nat. Comm.

Soppa, J., Kobayashi, K., Noirot-Gros, M.-F., Oesterhelt, D., Ehrlich, S.D., Dervyn, E., Ogasawara, N., and Moriya, S. (2002) Discovery of two novel families of proteins that are proposed to interact with prokaryotic SMC proteins, and characterization of the *Bacillus subtilis* family members ScpA and ScpB. Mol. Microbiol. 45: 59–71.

Sullivan, N.L., Marquis, K.A., and Rudner, D.Z. (2009) Recruitment of SMC by ParB-*parS* organizes the origin region and promotes efficient chromosome segregation. Cell 137: 697–707.

Tran, N.T., Laub, M.T., and Le, T.B.K. (2017) SMC progressively aligns chromosomal arms in *Caulobacter crescentus* but is antagonized by convergent transcription. Cell Rep 20: 2057–2071.

Treuner-Lange, A., Aguiluz, K., van der Does, C., Gomez-Santos, N., Harms, A., Schumacher, D., Lenz, P., Hoppert, M., Kahnt, J., Munoz-Dorado, J., and Søgaard-Andersen, L. (2013) PomZ, a ParA-like protein, regulates Z-ring formation and cell division in *Myxococcus xanthus*. Mol Microbiol 87: 235–253.

Tzeng, L., and Singer, M. (2005) DNA replication during sporulation in *Myxococcus xanthus* fruiting bodies. Proc. Natl. Acad. Sci. USA 102: 14428–14433.

Vallet-Gely, I., and Boccard, F. (2013) Chromosomal organization and segregation in *Pseudomonas aeruginosa*. PLoS Genet 9: e1003492.

Vasa, S., Lin, L., Shi, C., Habenstein, B., Riedel, D., Kühn, J., Thanbichler, M., and Lange, A. (2015) β-Helical architecture of cytoskeletal bactofilin filaments revealed by solid-state NMR. Proc. Natl. Acad. Sci. USA 112: E127–E136.

Vecchiarelli, A.G., Hwang, L.C., and Mizuuchi, K. (2013) Cell-free study of F plasmid partition provides evidence for cargo transport by a diffusion-ratchet mechanism. Proc. Natl. Acad. Sci. USA 110: E1390–E1397.

Viollier, P.H., Thanbichler, M., McGrath, P.T., West, L., Meewan, M., McAdams, H.H., and Shapiro, L. (2004) Rapid and sequential movement of individual chromosomal loci to specific subcellular locations during bacterial DNA replication. Proc Natl Acad Sci U S A 101: 9257–9262.

Wang, X., Brandao, H.B., Le, T.B., Laub, M.T., and Rudner, D.Z. (2017) *Bacillus subtilis* SMC complexes juxtapose chromosome arms as they travel from origin to terminus. Science 355: 524–527.

Wang, X., Le, T.B., Lajoie, B.R., Dekker, J., Laub, M.T., and Rudner, D.Z. (2015) Condensin promotes the juxtaposition of DNA flanking its loading site in *Bacillus subtilis*. Genes Dev 29: 1661–1675.

Wang, X., Liu, X., Possoz, C., and Sherratt, D.J. (2006) The two *Escherichia coli* chromosome arms locate to separate cell halves. Genes Dev. 20: 1727–1731.

Wang, X., Llopis, P.M., and Rudner, D.Z. (2013) Organization and segregation of bacterial chromosomes. Nat Rev Genet 14: 191–203.

Wang, X., Tang, Olive W., Riley, Eammon P., and Rudner, David Z. (2014) The SMC condensin complex is required for origin segregation in *Bacillus subtilis*. Curr. Biol. 24: 287–292.

Wu, S.S., Wu, J., and Kaiser, D. (1997) The *Myxococcus xanthus pilT* locus is required for social gliding motility although pili are still produced. Mol. Microbiol. 23: 109–121.

Yamaichi, Y., Bruckner, R., Ringgaard, S., Moll, A., Cameron, D.E., Briegel, A., Jensen, G.J., Davis, B.M., and Waldor, M.K. (2012) A multidomain hub anchors the chromosome segregation and chemotactic machinery to the bacterial pole. Genes Dev 26: 2348–2360.

Yamaichi, Y., Fogel, M.A., and Waldor, M.K. (2007) *par* genes and the pathology of chromosome loss in *Vibrio cholerae*. Proc. Natl. Acad. Sci. USA 104: 630–635.

Yatskevich, S., Rhodes, J., and Nasmyth, K. (2019) Organization of chromosomal DNA by SMC complexes. Ann. Rev. Genet. 53: 445–482.

Yu, W., Herbert, S., Graumann, P.L., and Gotz, F. (2010) Contribution of SMC (structural maintenance of chromosomes) and SpoIIIE to chromosome segregation in *Staphylococci*. J Bacteriol 192: 4067–4073.

