## Supplementary material for "SMC and the bactofilin/PadC scaffold have distinct yet redundant functions in chromosome segregation and organization in *Myxococcus xanthus*": All supplementary data

Department of Ecophysiology,

35043 Marburg, Germany


This file contains

- Supplementary Figure S1-S12
- Supplementary Table S1

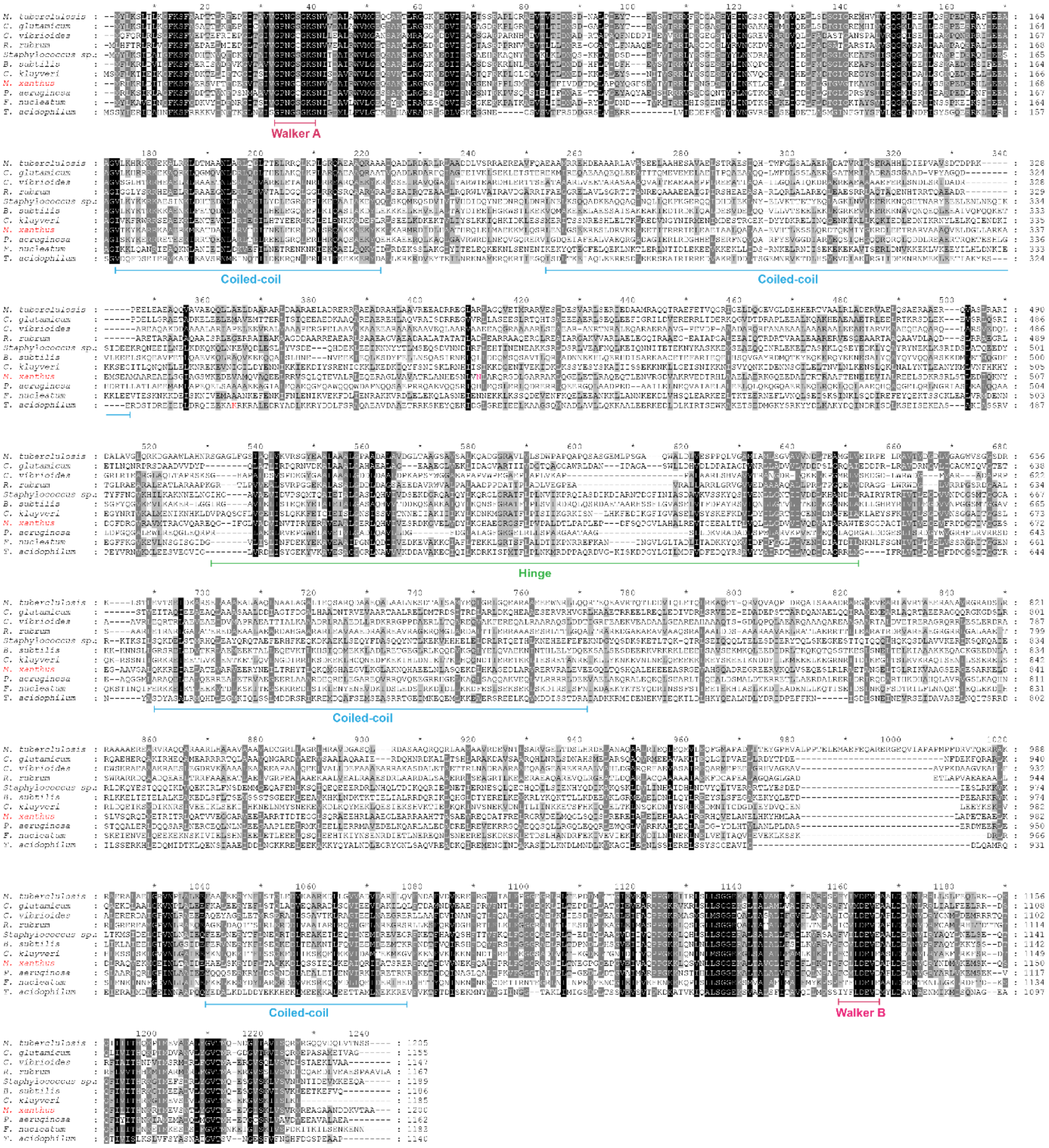


**Figure S1.** Sequence alignment of Smc proteins.
Alignment of *M. xanthus* Smc with Smc orthologs. White on black background indicates 100% conservation, white on dark grey background indicates 80% conservation, and black on light grey indicates 60% conservation. Conserved domains and sequence motifs are indicated. See also Fig. 1B.

**A**

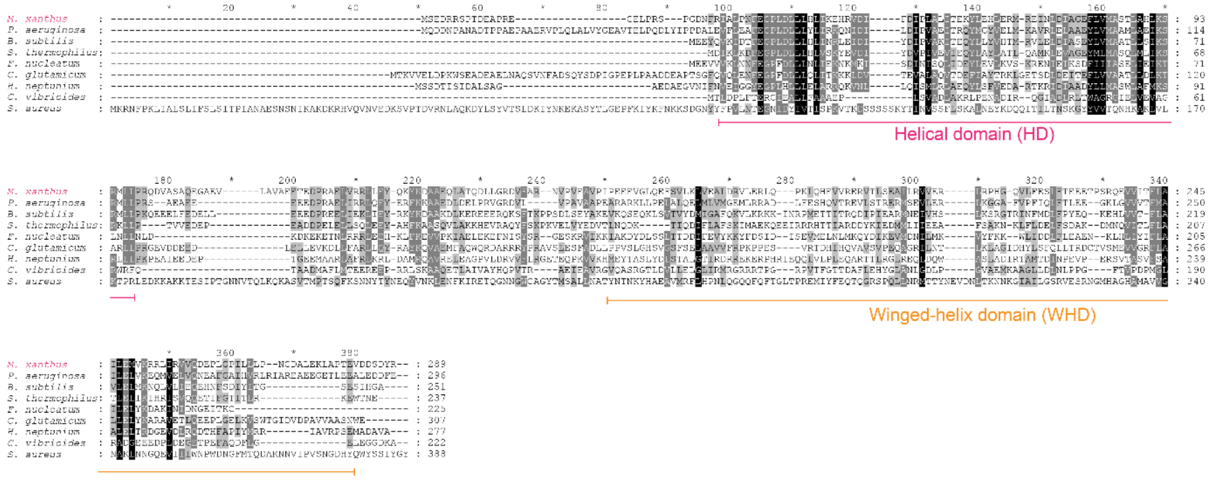


**B**

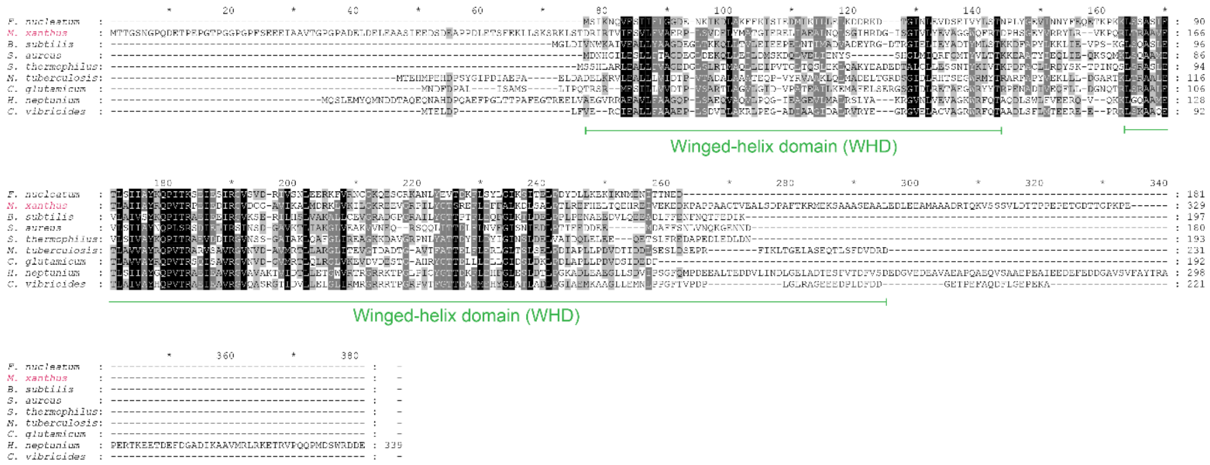


**Figure S2.** Sequence alignment of ScpA and ScpB proteins.
(A, B) Alignment of *M. xanthus* ScpA (A) and ScpB (B) with ScpA and ScpB orthologs. White on black background indicates 100% conservation, white on dark grey background indicates 80% conservation, and black on light grey indicates 60% conservation. Conserved domains are indicated. See also Fig. 1B.

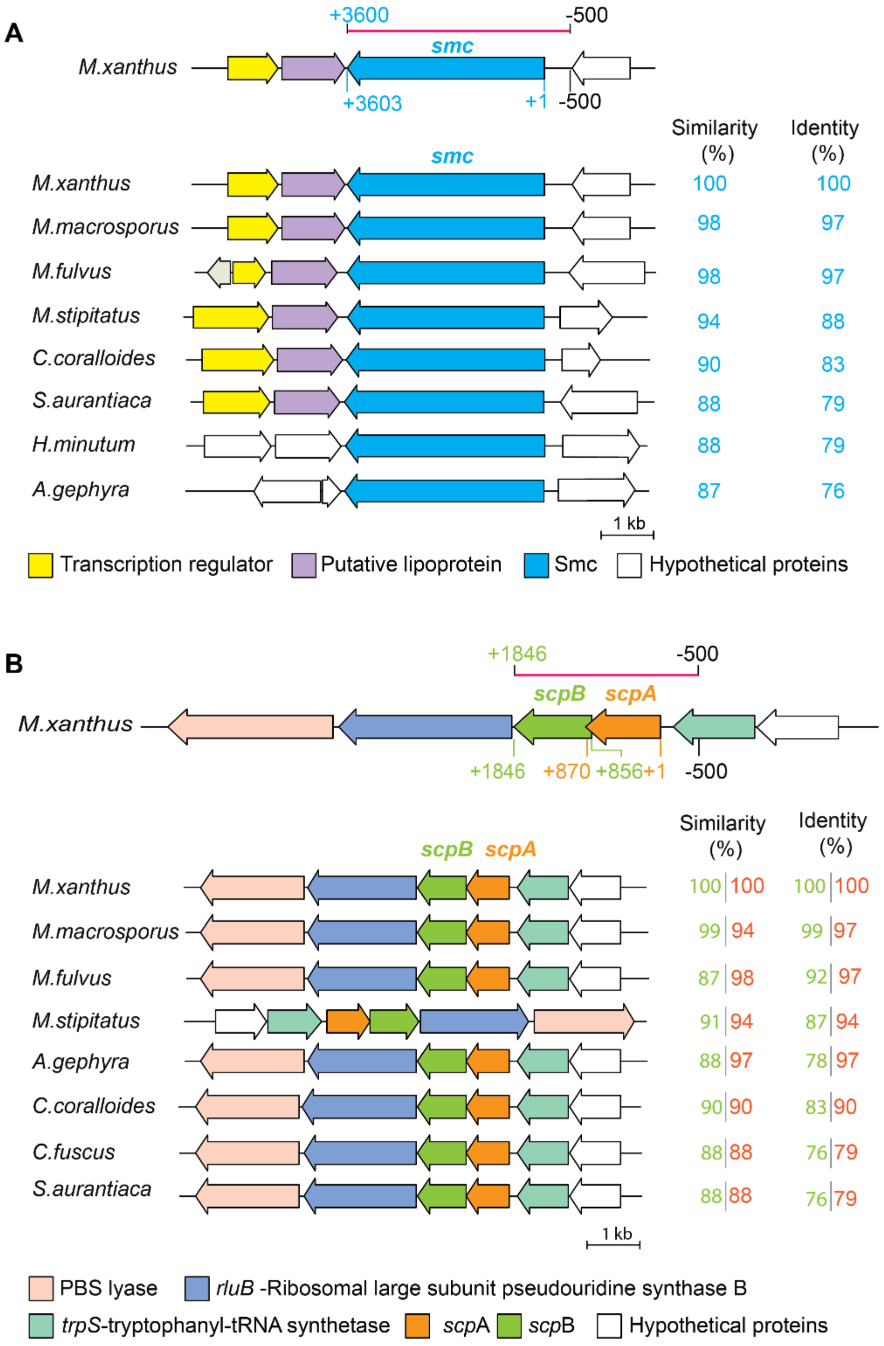


**Figure S3.** Genetic organization of *smc*, *scpA* and *scpB* genes in Myxococcales (A, B) Genetic context of *smc* (MXAN_4901) as well as *scpA* (MXAN_3841) and *scpB* (MXAN_3840) in *M. xanthus* and other fully sequenced Myxococcales genomes. Direction of arrows indicates 5' to 3' orientation of gene. Coordinates are shown relative to the first nucleotide of *smc* and *scpA,* respectively. % protein similarity and identity is shown on the right. The fragments used in complementation experiments are indicated by a red line above *smc* and *scpAB* and with the native promoter (Pnat) extending 500 bp upstream of the start codon of *smc* and *scpA*.

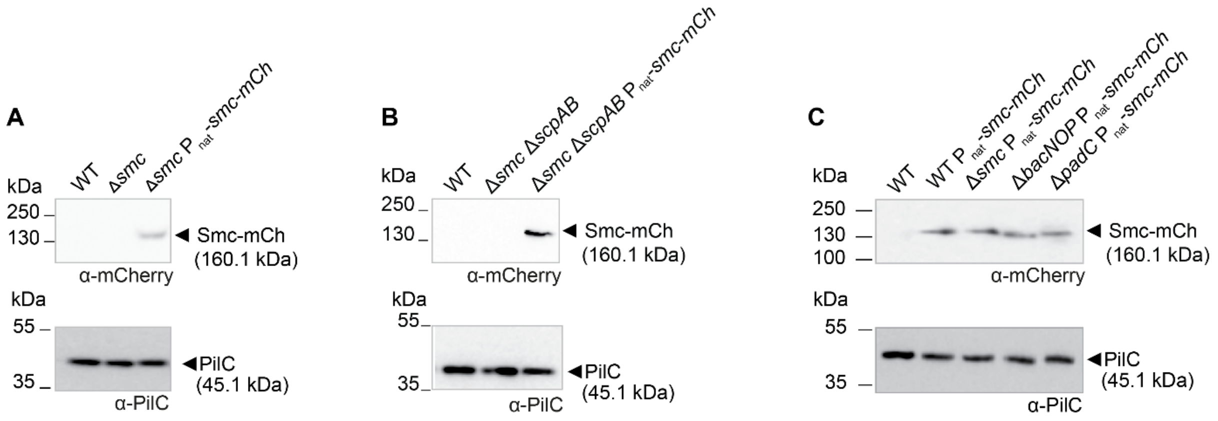


**Figure S4**. Immunoblot detection of Smc-mCherry accumulation.
(A, B, C) Accumulation of Smc-mCherry in strains of the indicated genotypes. Cells were grown at 25°C. Protein from an equal amount of cells was loaded per lane and probed with the indicated primary antibodies. The PilC immunoblot served as a loading control. Calculated molecular masses of Smc-mCherry and PilC are indicated. Molecular size markers are indicated on the left. In the complementation strains, *smc-mCherry* was expressed from its native promoter ($P_{nat}$) from a plasmid integrated in a single copy at the Mx8 *attB* site.

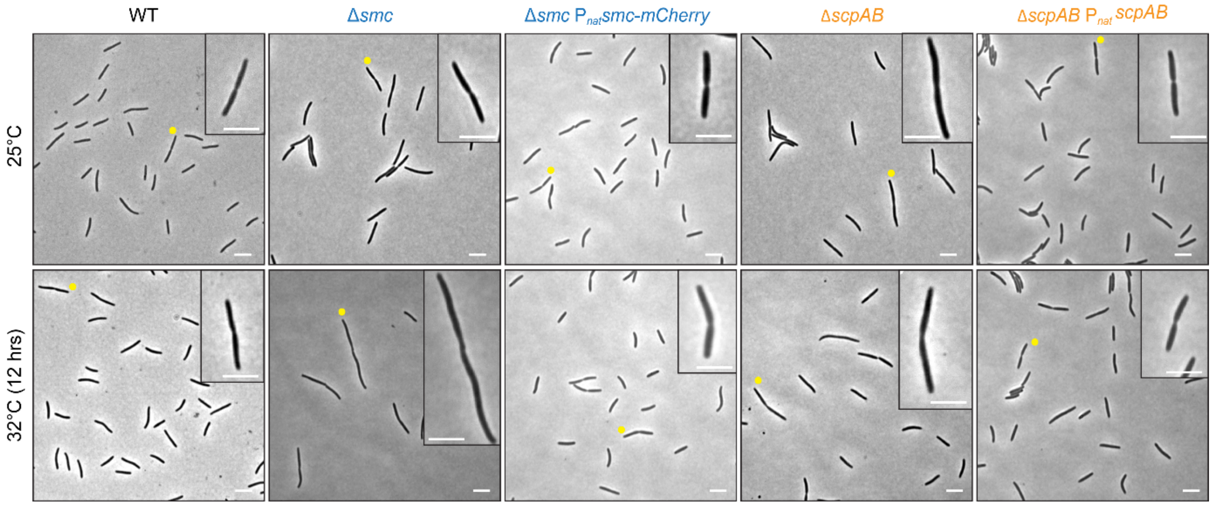


**Figure S5**. Morphology of cells lacking Smc or ScpAB
Phase contrast images of WT, Δ*smc*, Δ*scpAB* strains and the complementation strains grown at 25°C and 32°C (12 hrs). Yellow dots indicate cells with constriction shown in inset. Scale bars, 5 µm.

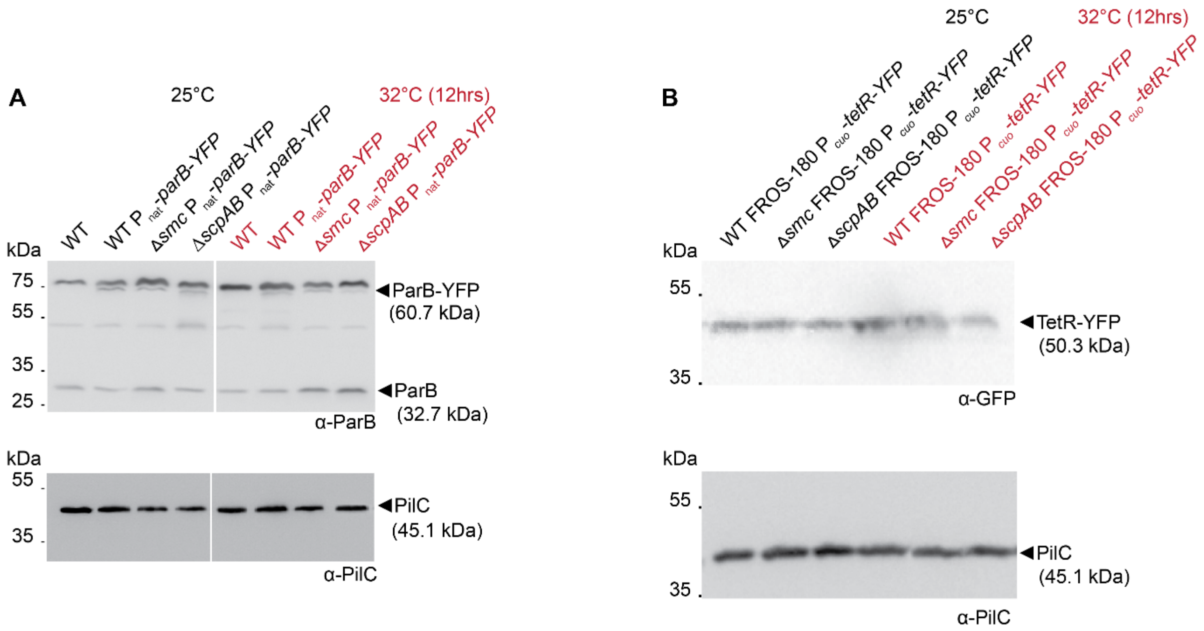


**Figure S6.** Immunoblot detection of ParB-YFP and TetR-YFP accumulation.
(A, B) Cells were grown at the indicated temperatures. Total protein from an equal amount of cells was loaded per lane and probed with the indicated primary antibodies. The PilC immunoblot served as a loading control. Calculated molecular masses of ParB-YFP, ParB, TetR-YFP and PilC are indicated. Molecular size markers are indicated on the left. *parB-YFP* was expressed from its native promoter ($P_{nat}$) from a plasmid integrated in a single copy at the Mx8 *attB* site and *tetR-YFP* was expressed from the cupper inducible $P_{cuo}$ from a plasmid integrated in a single copy at the Mx8 *attB* site. In B, cells were grown in the presence of 300 µM $CuSO_4$ for 6 hrs to induce expression of *tetR-YFP* from the $Cu^{2+}$ inducible $P_{cuo}$.

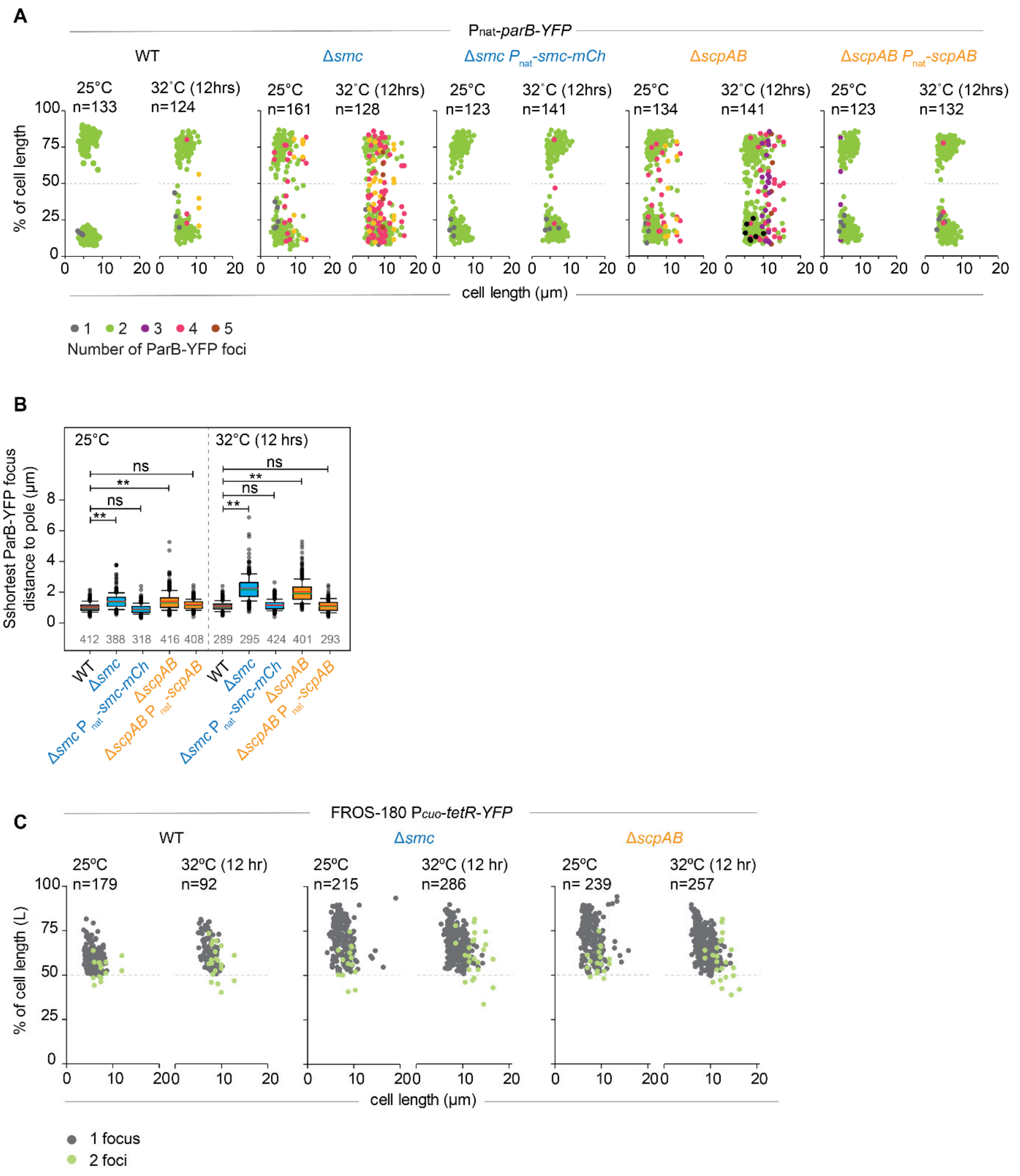


**Figure S7**. Analysis of ParB-YFP localization as well as TetR-YFP localization in FROS-180 strains

(A) Quantification of ParB-YFP localization pattern in strains of indicated genotypes at the indicated temperatures. Diagram indicates foci localization as % of cell length and as a function of cell length. The colour code indicates the number of foci per cell. N, number of cells analyzed from three biological replicates.

(B) Quantification of shortest distance between a pole and a ParB-YFP focus. Cells were treated as in A. Numbers below indicate number of cells from three biological replicates. Box plot is as in Fig. 2B. **, p <0.001; ns, not significant in Mann-Whitney test.

(C) Quantification of FROS-180 localization using TetR-YFP. Cells of the indicated genotypes were grown at the indicated temperatures. All strains contain the FROS-180 and P*cuo-tetR-YFP* constructs. Cells were grown in the presence of 300 µM $CuSO_4$ for 6hrs before microscopy. Diagram indicates foci localization as % of cell length and as a function of cell length. The colour code indicates the number of foci per cell. N, number of cells analyzed from three biological replicates.

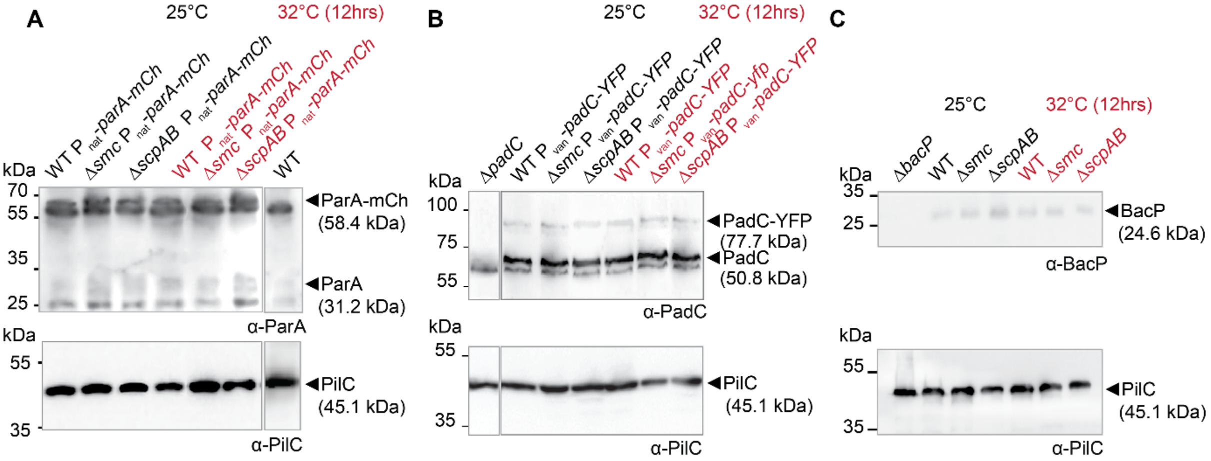


**Figure S8**. Immunoblot detection of ParA-mCherry, PadC-YFP and BacP accumulation
(A) Immunoblot detection of ParA and ParA-mCherry accumulation in strains of the indicated genotypes and grown at the indicate temperatures. Total protein from an equivalent number of cells was loaded per lane and probed with the indicated antibodies. The PilC immunoblot served as a loading control. *parA-mCherry* was expressed from its native promoter ($P_{nat}$) from a plasmid integrated at the *attB* site in merodiplod *parA*$^{+}$*/parA-mCherry* strains. All samples were separated on the same gel; the gap indicates lanes removed for presentation purposes. Calculated molecular masses of ParA-mCherry, ParA and PilC are indicated. Molecular size markers are indicated on the left.
(B) Immunoblot detection of PadC and PadC-YFP accumulation in strains of indicated genotypes and at indicated temperatures. Experimental details as in (A). *padC-YFP* was expressed from a vanillate-inducble promoter ($P_{van}$) from a plasmid integrated at the *MXAN_18-19* site in the presence of 5 μM vanillate in merodiploid *padC*$^{+}$*/padC-YFP* strains. All samples were separated on the same gel; the gap indicates lanes removed for presentation purposes. Calculated molecular masses of PadC-YFP. PadC and PilC are indicated. Molecular size markers are indicated on the left.
(C) Immunoblot detection of BacP accumulation in strains of indicated genotypes and at the indicated temperatures. Experimental details as in (A).

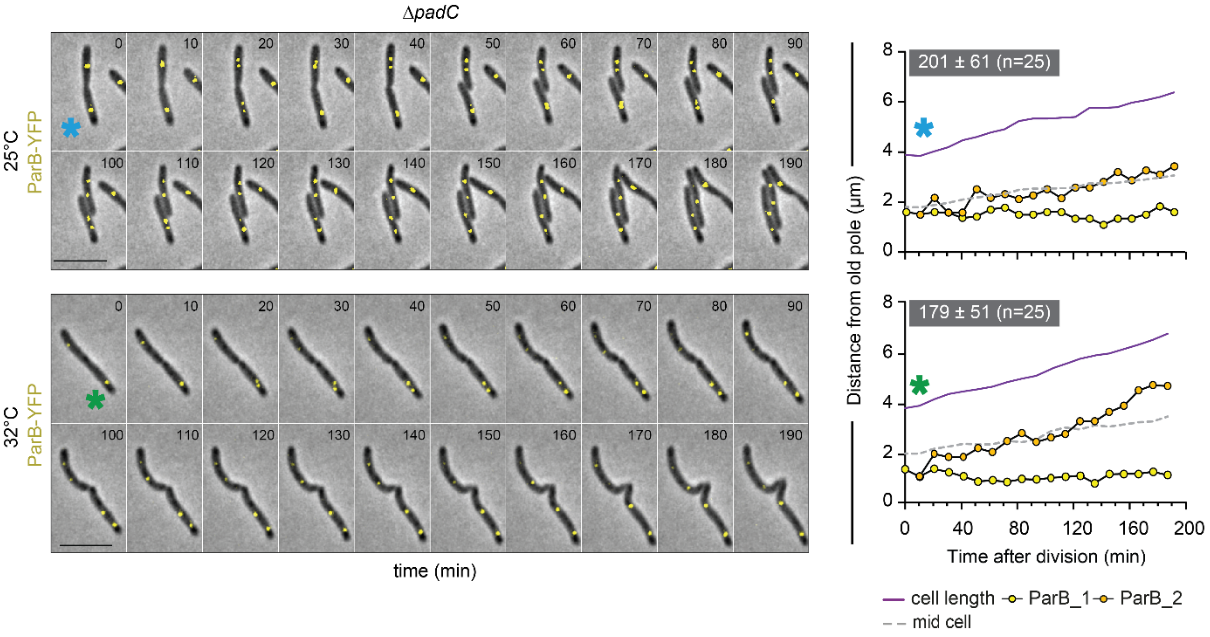


**Figure S9**. Lack of PadC causes a ParB·*parS* complex segregation defect.
Cells were grown at 25°C and transferred to a 1.0% agarose pad supplemented with 0.2% casitone and imaged after 15 min and then every 10 min at 25°C and 32°C. Montages (left panels) and line graphs of ParB-YFP trajectories (yellow, orange) (right panels) are shown. Coloured asterisks in montages indicate cells for which ParB-YFP trajectories are shown on the right. Scale bars, 5 µm. Average time ± SD for translocation of ParB-YFP focus from the old to the subpolar region at the new pole is indicated in white on the grey background together with the number of cells analysed. ParB-YFP was expressed ectopically from the native *parB* promoter in merodiploid *parB*$^+$/*parB-YFP* strains. Strain contained the Δ*mglA* mutation to inactivate the motility systems.

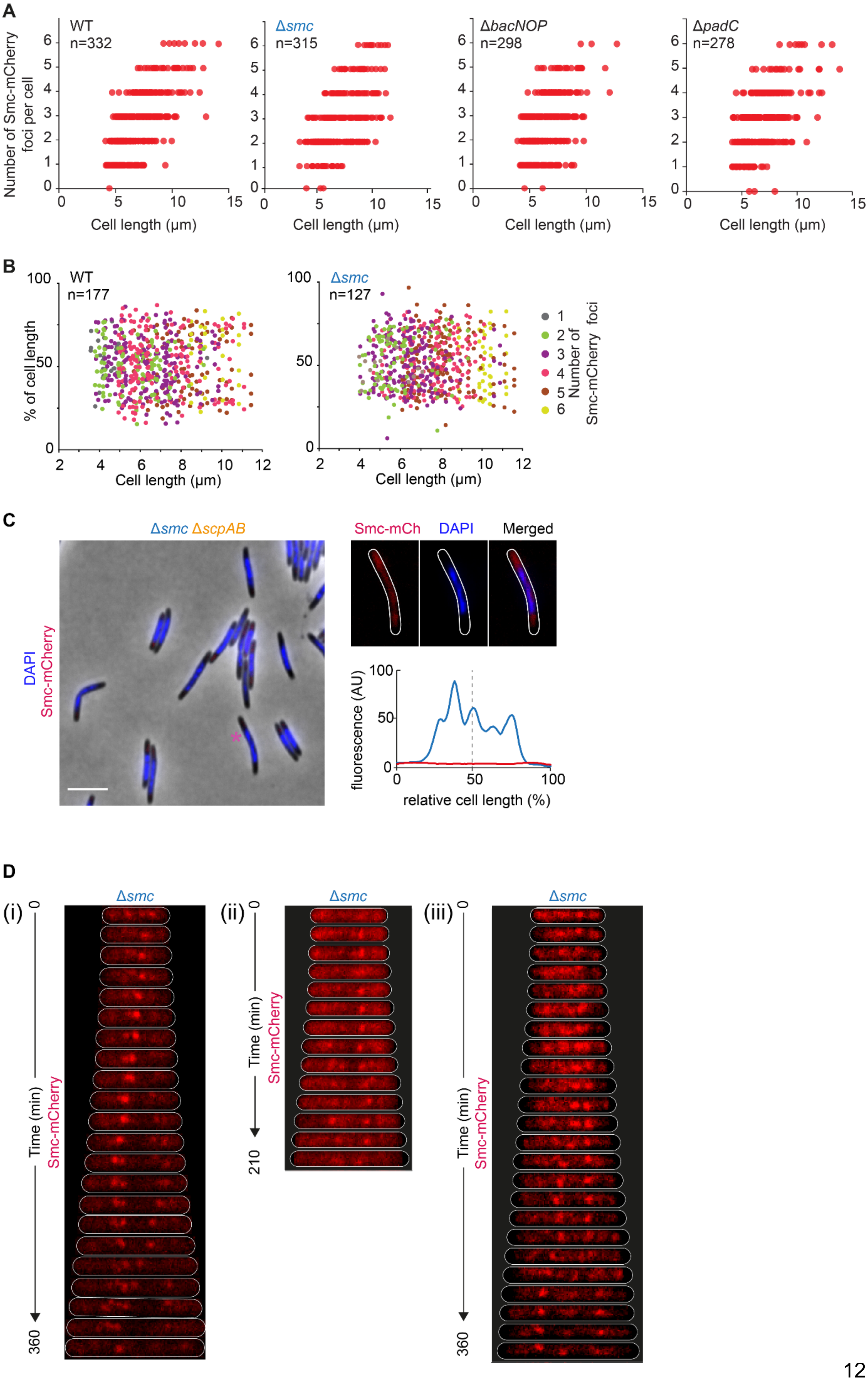
A
WT n=332
Δsmc n=315
ΔbacNOP n=298
ΔpadC n=278
Number of Smc-mCherry foci per cell
Cell length (µm)
B
WT n=177
Δsmc n=127
% of cell length
Cell length (µm)
Number of Smc-mCherry foci
C
Δsmc ΔscpAB
DAPI
Smc-mCherry
Smc-mCh
DAPI
Merged
fluorescence (AU)
relative cell length (%)
D
(i)
(ii)
(iii)
Δsmc
Time (min)
Smc-mCherry

**Figure S10.** Localization of Smc-mCherry.
(A) Analysis of number of Smc-mCherry foci per cell as a function of cell length. The number of Smc-mCherry foci per cell was quantified and plotted as a function of cell length. N, number of cells analysed from three biological replicates.
(B) Quantification of Smc-mCherry localization in strains of indicated genotype. Diagram indicates Smc-mCherry foci localization as % of cell length and as a function of cell length. N, number of cells analyzed from three biological replicates. Cells were grown at 32°C.
(C) Analysis of Smc-mCherry localization in strain of indicated genotype. Cells were stained with DAPI and imaged. Cells were grown at 25°C. Left image shows overlay of phase contrast, DAPI and Smc-mCherry signals; images on the right, a representative cell showing the signals for Smc-mCherry, DAPI and an overlay; the line scan below shows the DAPI (green) and Smc-mCherry (red) signals of the cell shown above plotted as a function of relative cell length.
(D) Time-lapse microscopy of Smc-mCherry dynamics. Images were captured at 32°C with 15 min intervals on 1% agarose buffered with TPM and 0.2% casitone. Cells were straightened, aligned and kymographs were generated as described in Methods. The three cells show examples of Smc-mCherry localization dynamics. The strain contained the Δ*mglA* mutation to inactivate the motility systems.

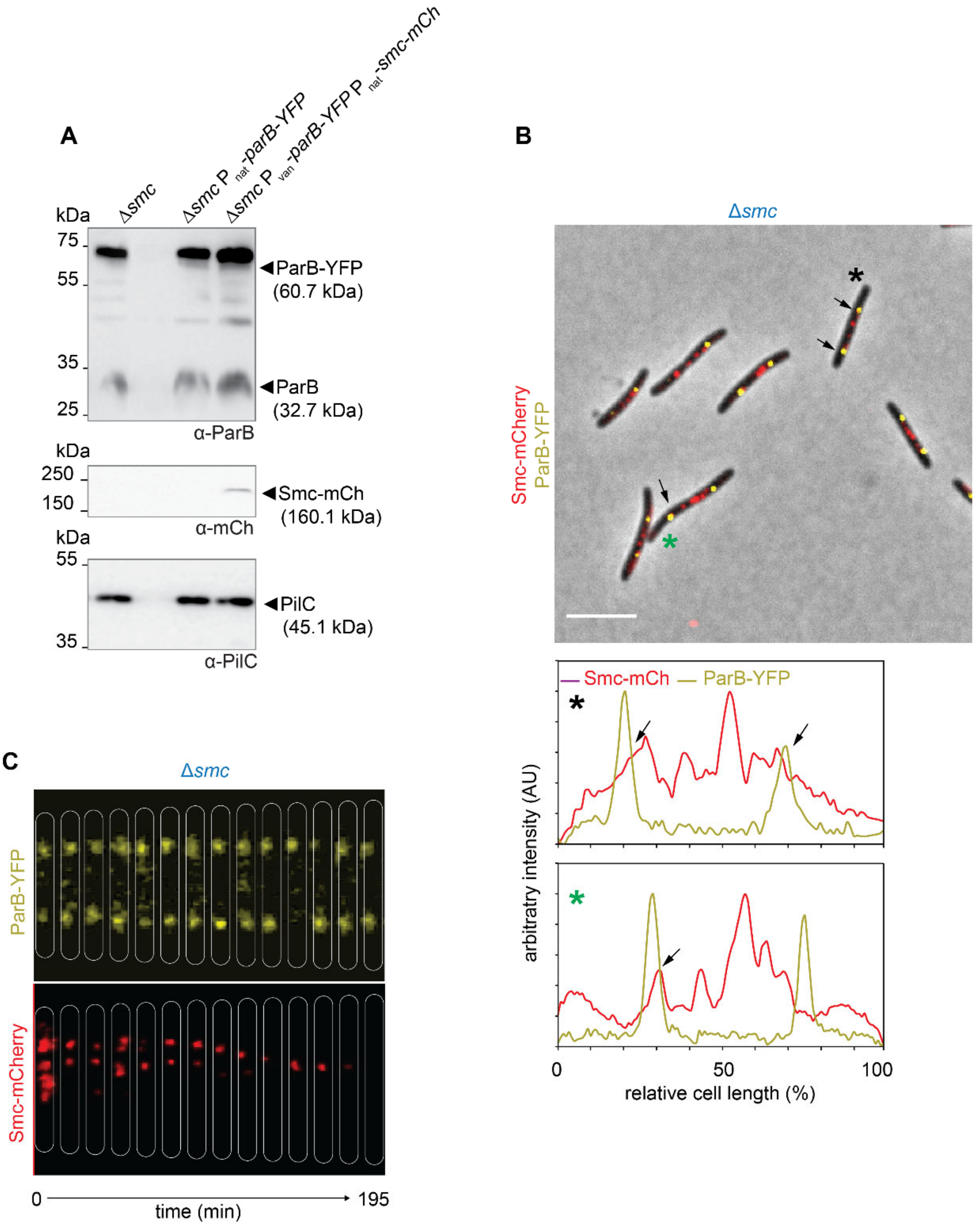


**Figure S11.** Colocalization experiments with Smc-mCherry and ParB-YFP.
(A) Accumulation of ParB-YFP and Smc-mCherry in strains of indicated genotypes. In the co-expression strain, ParB-YFP was expressed from the *attB* site under control of the vanillate promoter and Smc-mCherry was expressed under the control of its native promoter from the *MXAN_18-19* site. Cells were grown in the presence of 10 μM vanillate to induce synthesis of ParB-YFP for 6 hrs. The Δ*smc* mutant expressing ParB-YFP from its native promoter was used as a control for level of ParB-YFP accumulation after vanillate induction. Total protein from an equivalent number of cells was loaded per lane and probed with the indicated antibodies. PilC was used as a loading control.
(B) Colocalization experiments with Smc-mCherry and ParB-YFP
Cells were grown as in (A) and imaged. Image on top, overlay of phase contrast and Smc-mCherry (purple) and ParB-YFP (yellow) signals. Line scans below show fluorescence profile of Smc-mCherry and ParB-YFP signals as a function of relative cell length for the two

cells marked with asterisks in the image on top. Arrow indicates overlap of the two fluorescence signals.

(C) Time-lapse imaging of Smc-mCherry and ParB-YFP

Cells were grown as in (A) and imaged on 1% agarose with 10 µM vanillate and 0.2% casitone buffered with TPM. Images were captured at 15 min intervals. The cell was straightened, aligned and the kymograph generated as described in Methods. The strain contains a Δ*mglA* mutation to inactivate the two motility system.

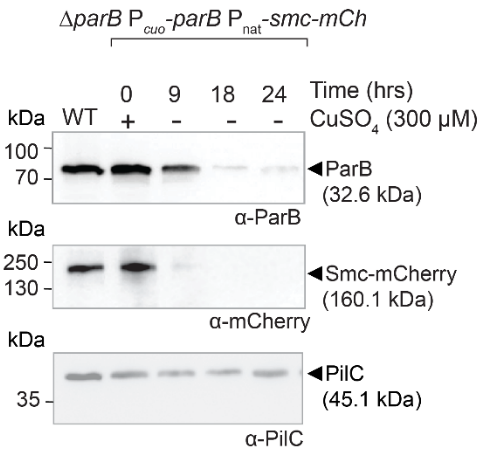


**Figure S12.** Depletion of ParB causes instability of Smc-mCherry.
Cells were grown in the presence of 300 μM $CuSO_4$; cells were transferred to copper-free medium at t=0 hrs and samples taken at the indicate time points. Total protein from an equivalent number of cells was loaded per lane and probed with the indicated primary antibodies. The PilC immunoblot served as a loading control. Calculated molecular masses of ParB, Smc-mCherry and PilC are indicated. Molecular size markers are indicated on the left.

**Supplementary Table 1.** Primers used in this study

| **Primer name** | **Sequence 5'-3'** [1] | **Comment** |
|---|---|---|
| Mxan-4901 A fwd | CTTCGCTGCGGGCGACCCGCG | deletion of *smc* (MXAN_4901) |
| Mxan-4901 B rev | CAAGCGGTTGGACATCACCGGC | |
| Mxan-4901 C fwd | GCCAACGACGACAAGGTGACG | |
| Mxan-4901 D rev | GACGCGGCGTCCGGCGCGGAC | |
| Mxan-4901Xba1fwd I | GCGTCTAGAATGCGAATCAAGCGGTTGGAC | |
| Mxan-4901EcoR1fwd J | GCGGAATTCTCCGCCGAGGGCATGTCAGAC | |
| Mxan-4901 BamH1 rev K | GCGGGATCCGGCGGAGCCCGCCGCCGTCACCTTGTC | |
| Mxan-4901 HindIII rev | GCGAAGCTTCTACGCCGCCGTCACCTTGTC | |
| Mx_scpAup-A- BamHI | GGAATTCGCCGGTGGACGCGCCCAAG | deletion of *scpAB* (MXAN_3841-3840) |
| Mx_scpAup-B-XbaI | GCTCTAGACGGATGTCCTGGATGGCGTCAAC | |
| Mx_scpAdw-C-XbaI | GCTCTAGAGTGACTACCGGTAGCAACGGACC | |
| Mx_scpAdw-D-HindII | CCCAAGCTTGTCTCCACCGCCGCCCGGGT | |
| Mx_scpBup-A | GGAATTCGGACCCACGCGCGGAACTG | |
| Mx_scpBup-B | GCTCTAGATGTCGTCGACCTCCGTGGG | |
| Mx_scpBdw-C | GCTCTAGACGGGCAGCATGGAAGAAGGAT | |
| Mx_scpBdw-D | CCCAAGCTTTCGAGCCGCACGCCGCC | |
| Mx_scpBup-A2 | GGAATTCAACTGGTGCGGCGCCTGCTGGA | |
| Mx_scpBup-B2 | GCTCTAGACTACCGGTAGTCACTGTCGTCG | |
| ParB-NdeI-F | GGAATTCCATATGGTGGTGAAAGCAGACATGCAGAA | Cloning |
| YFP-EcoRI-R | CGGAATTCTCACTTGTACAGCTCGTCCATGCCGAG | |

[1] Nucleotides in red indicate restriction sites used for cloning.
